# Brain network connectivity underlying remission in early psychosis: a whole brain model approach

**DOI:** 10.1101/2023.11.28.568844

**Authors:** Ludovica Mana, Ane López-González, Yasser Alemán-Gómez, Philipp S. Baumann, Raoul Jenni, Luis Alameda, Lilith Abrahamyan Empson, Paul Klauser, Philippe Conus, Patric Hagmann, Manel Vila-Vidal, Gustavo Deco

## Abstract

**Background:** Alterations in brain connectivity occur early during psychosis and underlie the clinical manifestations of the illness as well as patient functioning and outcome. After a first episode of psychosis (FEP), different trajectories are possible and best described by the clinical-staging model that places the patient along a continuum of conditions: from non-remitting chronic symptoms to full-remission, often followed by relapses. However, little is known about the differences in brain connectivity that could underlie these differences in clinical outcome.

**Methods:** In this study, we included resting-state fMRI and DSI data from a cohort of 128 healthy controls (HC) and 88 patients with early psychosis (EP) stratified based on their ability to remit after the FEP. In particular we focused on differences between stage IIIb,c remitting-relapsing (EP3R) and stage IIIa non-remitting (EP3NR) patients. We investigated alterations in resting-state functional connectivity (FC), and combined information derived from fMRI and DSI into generative whole-brain models of each condition to explore the underlying mechanisms.

**Results:** Opposite alterations in FC could be found in patients as compared to HC, depending on their stage. In non-remitting patients (EP3NR), we observed a reduction of FC, aligned with the reduced structural connectivity found in previous studies, while remitting-relapsing patients (EP3R) showed increased FC, potentially indicating a relevant compensatory mechanism. By means of a whole-brain network model, we showed that in HC a subset of areas is characterized by increased stability to prevent an oversynchronisation of the network, while in EP3 patients such property is lost. This alteration was more relevant in the EP3R than in EP3NR patients, probably indicating a compensatory response to the reduced effective conductivity (global coupling) highlighted by the model in both EP3 conditions as compared to controls.

**Conclusions:** These findings highlight the significance of categorizing patients into subgroups based on the progression of their psychotic disorders, providing insights into the factors contributing to heterogeneity in functional alterations. They enhance our understanding of the interplay between structural and functional properties, shedding light on the mechanisms of psychosis emergence, remission and progression, with potential implications for future therapeutic advancements.

## 1. Introduction

Clinical presentation and trajectory in early-stages psychotic disorders can vary widely among patients. Following a first episode of psychosis (FEP), approximately one-third of patients achieve full recovery ^1,2^, while the remainder progress to chronic illness, characterized by either a lack of remission, partial remission, or a cycle of complete remission followed by relapses ^3^.

From a clinical perspective, it is extremely important to better discriminate into different groups of patients based on clinical trajectory. To this aim, clinical staging –introduced to psychiatry by Fava and Kellner ^4^, and applied to psychotic disorders by McGorry and colleagues– is a fundamental tool ^5–9^. Additionally, mapping neurobiological markers, such as brain imaging features, onto clinical stages could further allow us to validate the boundaries of the clinical groups, broadening our understanding of psychotic disorder pathophysiology ^10,11^.

To date, most studies on early psychosis have primarily focused on discriminating patients with the aim of predicting their long-term recovery prospects following a first episode of psychosis, crucial for early intervention ^12–17^. However, from the neural perspective, equally important is understanding the distinctions among patients based on their ability to achieve short-term remission from FEP, regardless of prognosis. In fact, within the non-recovering patients, during the first years of treatment following a FEP, a noteworthy differentiation emerges between two distinct clinical profiles: those experiencing periods of complete remission between relapses and those who either do not remit or achieve only partial remission ^5^. This distinction holds the key to comprehending the neural mechanisms enabling a subset of patients to effectively overcome the acute symptomatic crisis and temporarily restore healthy functioning ^18^. Delving deeply into this phenomenon is crucial, as it could unveil compensatory mechanisms and might facilitate the identification of biomarkers associated with clinical staging, providing valuable insights into the mechanisms underlying disease progression. Moreover, it could help explain the heterogeneity of brain alterations in the literature. In fact, while studies investigating anatomical alterations in psychotic disorders are fairly coherent in reporting decreased structural connectivity (SC) ^19–22^, the way such changes translate into functional alteration remains unclear ^23^. Findings on functional alterations are indeed less consistent ^24,25^, with a significant portion of studies describing reductions in functional connectivity (FC), and others reporting increases ^26–29^.

However, neurofunctional differences between remitting-relapsing and non-remitting patients have been scarcely studied and underlying correlates remain unclear. Moreover, we still lack a thorough comprehension of how local alterations translate into global network changes, and how these in turn translate into clinical manifestations.

The purpose of this study is to address these gaps by employing connectivity analyses and computational approaches to explore the neural processes that enable a subgroup of patients to compensate for the alterations that lead to psychotic episodes and to achieve temporary remission, in contrast to other patients who fail to do so. Specifically, we aim to disentangle the underlying functional compensation mechanism into global and local components given the structural properties of the network.

To this aim, we included resting-state functional magnetic resonance imaging (fMRI) and diffusion spectrum MRI (DSI) data from a cohort of 128 healthy controls and 88 patients with early psychosis (within the first 3 years following a treated FEP in an early intervention service) stratified into distinct groups based on their clinical stage. Patients were classified into stage II (first-episode), stage IIIb,c remitting-relapsing (remission with one or more relapse), and stage IIIa non-remitting (incomplete remission), according to the course of the illness until the time of scanning ^5^. While a previous study on an overlapping sample of the same dataset showed a progressive decrease in structural connectivity across stages, it didn’t explore differences in functional connectivity or distinguish between remitting-relapsing and non-remitting patients ^30^. This current work focuses specifically on these questions, examining whether global and local FC measures significantly differ between remitting-relapsing and non-remitting patients compared to healthy controls. Moreover, we combined information derived from both fMRI and DSI to build a generative whole-brain model that could optimally fit the empirical data of the controls. Complementing it with an additional toy-model, we used it to extract hidden properties of the network that cannot be directly measured. This approach allowed us to investigate how brain dynamics interact with connectomes in the healthy brain. Finally, we built models to explain the empirical data of each subgroup of patients, investigating potential damaging and compensatory mechanisms underlying their pathological alterations.

## 2. Materials and methods

### 2.1. Participants

We included 216 subjects (88 EP patients, 128 HC) aged 18-35 years [26.3 ± 0.6]. EP patients were recruited from the Treatment and Early Intervention in Psychosis (Tipp) program of the Lausanne University Hospital, Switzerland ^31^, meeting psychosis threshold and specific criteria (see supplementary materials). HC were age, gender, and handedness-matched, without mood or substance use disorders, and no family history of psychosis. Exclusion criteria included neurological disorder, severe head trauma, or mental disability (IQ<70). Informed written consent was obtained, and the study was approved by the local ethics Committee.

### 2.2. Diagnosis, clinical variables, and staging

A consensus diagnosis procedure was elaborated in the framework of the TIPP program and is described in supplementary material. The clinical stage was rated as the highest stage achieved at the time of imaging, therefore cross-sectionally. The patients were initially stratified into four distinct groups (**Fig. 1**) based on a consensus assessment by two experienced psychiatrists, according to the clinical staging model proposed by McGorry et al. 2006.

**Fig.1:**
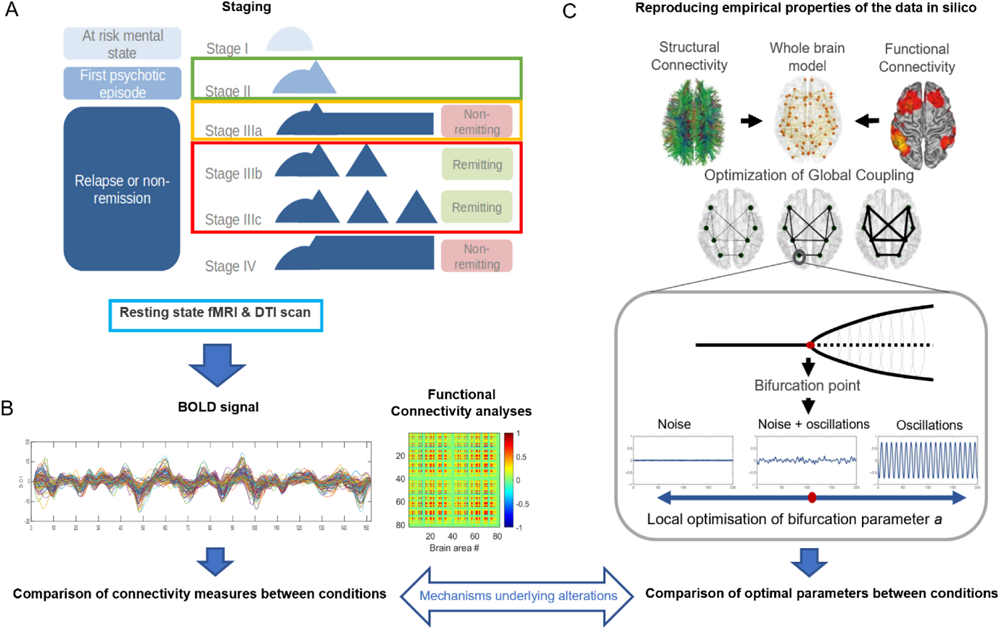
Schematic representation of the analysis workflow, featuring clinical staging model, empirical connectivity analyses and whole-brain model fitting. *(A) Stage I: early or late prodromal patients with mild or subthreshold symptoms; Stage II: first-episode of psychosis (i.e., ‘discrete disorder’); Stage IIIa: incomplete remission; Stage IIIb: one relapse; Stage IIIc: multiple relapses; Stage IV: chronic outcome with severe, persistent illness. This study includes resting state fMRI data from patients with early psychosis (EP) classified in stage 2, stage3 non-remitting (IIIa), stage3 remitting-relapsing (IIIb or IIIc), as well as from healthy controls (HC). Figure adapted from* ^30^*, Copyright © 2019 Griffa et al., under CC BY 4.0. (B) Functional connectivity (FC) matrices were obtained for each subject by computing the Pearson correlation of the BOLD signals between each pair of nodes across time. Global and local measures of connectivity were computed to investigate significant alterations between condition. (**C**) Structural and functional information from the empirical data was used to build a whole-brain dynamical model and fit it to reproduce the empirical FC features of the HC and the different EP groups independently in order to investigate the mechanisms underlying pathological alterations. Figure adapted from Deco et al., 2019, Copyright © 2017 Deco et al., under CC BY 4.0*.

Stage II (EP2) included first-episode psychosis patients with one episode. Stage III (EP3) included three subgroups: IIIa (incomplete remission), IIIb (relapse), and IIIc (multiple relapses). Due to the study’s goal of investigating neural mechanisms in remission, and to the elevated heterogeneity of EP2 patients, we focused on EP3 patients. Stage IIIb and IIIc were aggregated in a remitting-relapsing (EP3R) subgroup with 31 patients, while stage IIIa formed the non-remitting (EP3NR) subgroup with 20 patients. EP2 patients, being a more diverse group with various recovery durations and outcomes, were not included in the primary analysis, but their results were examined separately in the supplementary materials for completeness. Additional information can be found in supplementary materials.

### 2.3. Neuroimaging data acquisition and processing

MRI data was acquired from two 3-Tesla scanners, the Magnetom TrioTim and PRISMA from Siemens Medical Solutions. The MRI sessions included a T1-weighted MPRAGE sequence (TR = 2300 ms, voxel size = 1 x 1 x 1.2 mm³), a diffusion spectrum imaging (DSI) sequence with 1 b0 acquisition and 128 diffusion weighted directions (TR = 5900 ms, voxel size = 2.2 x 2.2 x 3 mm³), and a resting-state functional MRI (rs-fMRI) sequence with TR = 1920 ms, and 3.3mm isotropic. The acquisition times for the MPRAGE-T1w, DSI and rs-fMRI sequences were approximately 7, 13 and 9 minutes. Image quality assessment, including visual inspection and quality control metrics, ensured dataset quality. More details can be found in a previous work^32^ and in supplementary materials.

Image processing involved grey matter parcellation with skull-stripping using CAT12 and FreeSurfer for cortical surface reconstruction and subcortical parcellation. These regions were combined with the hippocampus subfields,^33^ the thalamic nuclei^34^ and a parcellation of the brainstem ^35^ to create a final subdivision of grey matter in 115 regions of interest. For DSI preprocessing, fiber tracking and structural connectivity computation, the images were processed using the image processing packages Mrtrix3 and FSL. rs-fMRI preprocessing was conducted with fMRIPrep, including motion correction, co-registration with anatomical images, spatial normalization, slice-timing correction, smoothing, and nuisance signal regression. Neuroimaging procedures are described in full detail in supplementary materials.

### 2.4. Measures of connectivity

The structural connectivity (SC) between each pair of grey matter regions of interest was quantified as the density of streamlines connecting the two regions, accounting for the differences in region sizes. This resulted in 115-nodes, weighted undirected brain-networks. To ensure consistency and minimize biases, connections present in less than 50% of subjects (both controls and patients) were excluded.

Functional connectivity (FC) was measured as the Pearson correlation between filtered BOLD signals (0.04-0.07 Hz) ^36^ of each pair of brain areas over the entire recording period, resulting in a 115x115 undirected matrix per subject. To validate our findings, we repeated all analyses using an independent FC measure -instantaneous phase coherence (iFC)-, which assesses the consistency of phase relationships across time. Node strength (*FC_sj_*) were computed as the sum over columns of the pairwise FC matrix and represents the overall level of connectivity of each individual node *i*. Significant differences in group averaged pairwise connectivity and in node strength were investigated between each EP group (EP3NR, EP3R) and HC. Next, we evaluated whether there were consistent changes in functional strength across all nodes between each EP group and HCs by comparing group average strength (*FCS_j_^condition^*) distributions. Finally, we compared the mean difference of strength in each area between the pathological conditions and HCs:

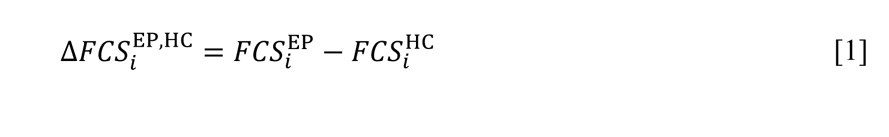

and tested whether the obtained global difference of strength between conditions was large enough to reject the null hypothesis that the two conditions had the same mean value (H0: Δ=0). Moreover, global network-level analysis involved calculating overall level of coupling through mean FC values, and computing root-mean-square signal amplitude, segregation, and integration. Wilcoxon ranksum test was used to investigate differences between groups. A threshold of α = 0.05 was applied.

We additionally explored the relationship between structural and functional connectivity by computing the Pearson correlation between the upper triangular elements of the FC and SC matrixes, as well as between global values of FC and SC in each condition. All connectivity analyses are described in full detail in supplementary materials.

### 2.5. Modelling whole-brain dynamics in patient groups and controls

For each condition, we created a whole-brain model using structural and functional connectivity data, adjusting parameters to replicate empirical data (**Fig. 1C**), as described in supplementary materials. The model’s differential equations describe how local and global brain dynamics are influenced by structural connectivity, using the normal form of a supercritical Hopf bifurcation:

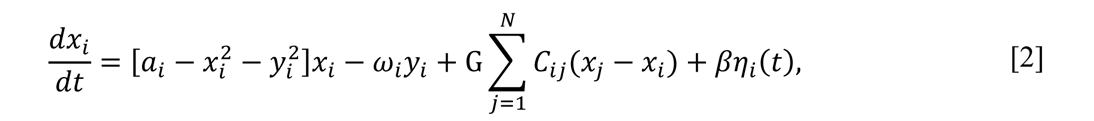

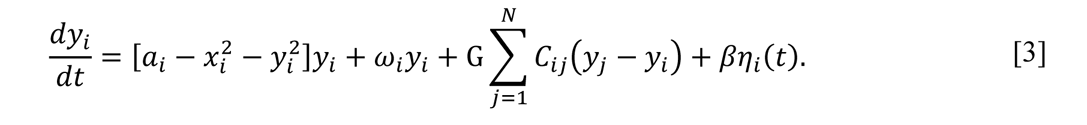

The connectivity between the different nodes was set to be equal to the empirical anatomical structural connectivity matrix *C* = (*C*_*ij*_) derived from corresponding DSI (N= 115 areas) for each condition, and the strength of these connections was scaled by the global coupling parameter *G*, which is believed to reflect the effectiveness of conductivity of the structural connections. The local dynamics were represented by bifurcation parameters (*a_i_*). According to this model, a bifurcation occurs at *a_i_*= 0 (edge of criticality), so that when *a_i_* assumes negative values (subcriticality) the activity of the node is described as noise, while when it assumes positive values (supercriticality), the behaviour becomes oscillatory with an intrinsic frequency determined by the parameter *f_i_* = *ω*_*i*_/2*π*. In the current work, we set the frequency within the 0.04-0.07 Hz range, and we derived it from the empirical data, as given by the averaged peak frequency of the narrowband BOLD signals of each brain region.

We optimized *G* and *a_i_* iteratively, and we ran 300 simulations for each parameter combination. We assessed model goodness of fit using functional connectivity (FC) and functional connectivity dynamics (FCD) and selected the combination of parameters for which the mean Kolmogorov-Smirnov (KS) distance was minimised as the optimal working point of the model. Next, we compared optimal global and local parameters between conditions.

#### 2.5.1. Additional statistical analyses

We then explored the relation of optimal values of *a_i_* with functional and structural node strengths in HC, by means of a generalised linear model. We also investigated how the change in the optimal value of bifurcation parameters in each pathological condition as compared to HC (Δ*α*^EP,HC^) relates with the corresponding values of *FCS*_*i*_^HC^ and *SCS*_*i*_^HC^. Additionally, we investigated how changes in optimal *a_i_* (Δ*α*_*i*_^EP,HC^) correlated with changes in empirical FC strength in pathological conditions (Δ*FCS*_*i*_^EP,HC^). More detailed description of the methods can be found in supplementary materials.

### 2.6. The impact of local alterations on the network dynamics

Additionally, we aimed to explore how local bifurcation parameters influenced global network dynamics by constructing a simplified network of coupled Hopf oscillators (two communities connected through a single node) and altering the bifurcation parameters of nodes, examining their impact on both spontaneous dynamics and responses to external stimuli (**Fig. 4**).

## 3. Results

### 3.1. Subjects and clinical staging

We investigated alterations in fMRI data among young adults in the early phases of psychosis (EPPs) classified into stage II (37 subjects), stage III remitting-relapsing (31 subjects), and stage III non-remitting (20 subject) subgroups, primarily focusing on exploring differences between the subgroups within stage III and comparing them with healthy controls (HCs).

**Supplementary Table 1** provides details on the demographics and clinical scores, and **Supplementary Table 2** the corresponding statistical comparisons computed through unpaired t-test. No significant differences in age, gender, or handedness were observed between EPPs and HCs. Stage III patients were, on average, older and had significantly longer durations of illness compared to stage II patients. It is important to acknowledge that the differences in the duration of illness between groups are inherent to the clinical-staging criteria used in this study. No difference in age or duration of illness was instead observed within stage III between remitting-relapsing (EP3R) and non-remitting patients (EP3NR), reducing confounding variables and making them more directly comparable.

Overall, EPPs had significant lower GAF scores compared to the HCs. Among patients, stage II and stage III showed no significant differences in GAF scores, PANSS scores, or medication doses at the time of fMRI. However, within stage III, EP3NR patients had higher PANSS general and total scores and lower GAF values than EP3R. PANSS negative and positive scores were also notably higher in EP3NR, though not statistically significant. No difference in medication dose was found between EP3R and EP3NR patients. The difference in symptom severity at the time of scanning serves as the basis for this alternative subdivision (EP3R vs EP3NR). It allows us to investigate neurological distinctions between two groups with equivalent disease progression stages (age, illness duration, and medication level) but significantly different clinical manifestations.

### 3.2. Empirical functional connectivity alterations (FC)

The first step in our analysis consisted of searching for alterations in network synchronisation between conditions (HC, EP3R, EP3NR), as measured by the FC measures computed from pairwise coupling of BOLD signals between all regions of interest.

#### Global network-level differences

No significant differences in global functional connectivity, integration, or mean amplitude were observed between conditions. However, trends indicated that remitting-relapsing patients tended toward increased connectivity, while non-remitting patients leaned toward decreased values compared to healthy controls **(Supplementary Fig. S1).**

#### Pairwise connectivity differences

We calculated functional connectivity differences between each EP group (EP3R, EP3NR) and HCs for each pair of brain areas. **Fig. 2A** shows the difference matrices. To enhance clarity, we only display pairs with p<0.01 significance, masking others. No differences survived multiple comparison correction, but distinct condition-specific trends emerged. EP3R patients exhibited increased connectivity in the majority of altered pairs, while EP3NR patients showed decreased connectivity in most of affected pairs.

**Fig.2:**
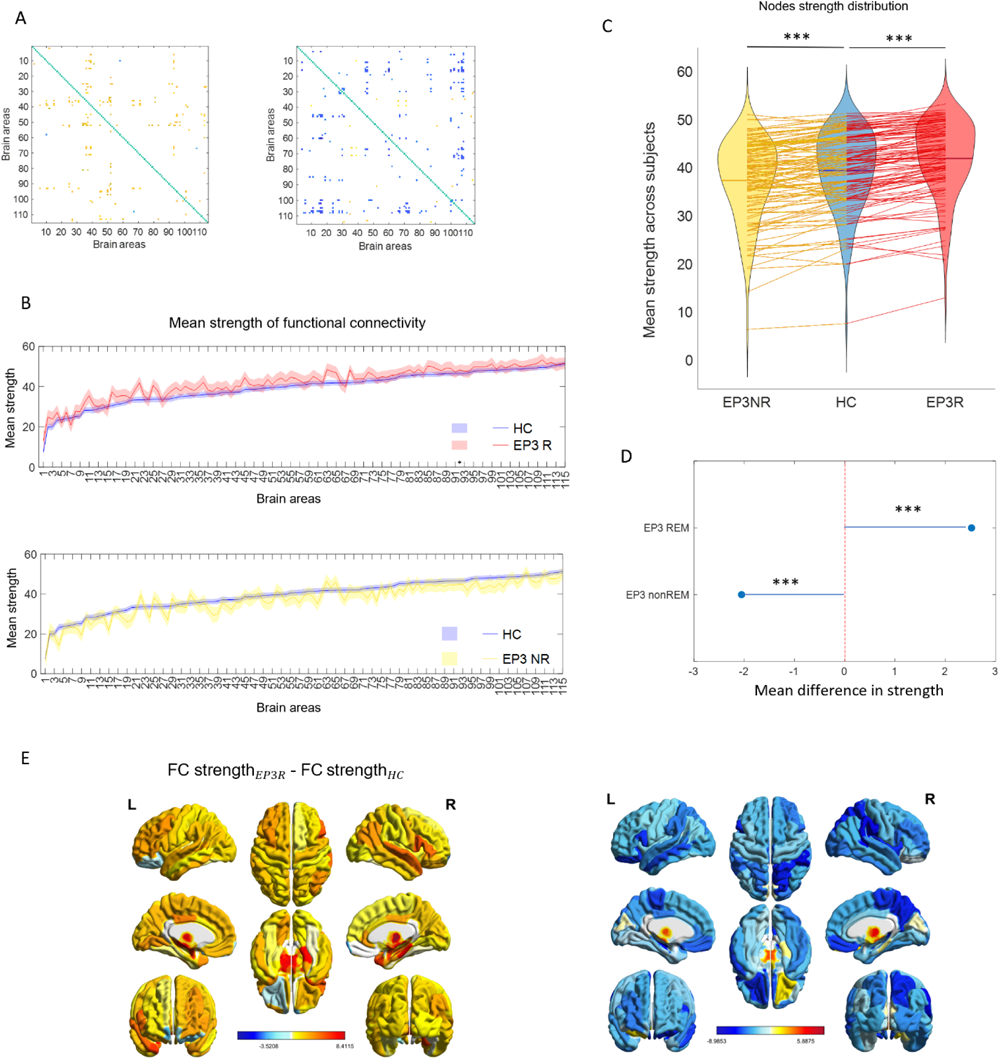
Empirical functional connectivity alterations: **(A)** Differences in group mean pairwise functional connectivity for each pair of brain areas between the pathological conditions and controls. Only the areas with a significant difference with p<0.01 are shown, while all the other areas have been masked. No significance survived multiple comparison correction. **(B)** Group mean connectivity strength for each of the 115 regions, ordered by mean regional strength in healthy controls. Solid lines and shaded areas represent the median and the standard error across subjects, respectively. None of the comparisons survived multiple comparison analysis in any of the conditions. Both in (A) and (B) it is possible to notice opposite trends of alteration in the two condition as compared to controls (increased FC in EP3R vs decreased FC in EP3NR). **(C)** Distributions of mean FC strength across subjects in pathological and healthy condition. Lines connect the value of mean strength in each area in healthy condition (central, blue) to the value of strength in the corresponding area in the two pathological conditions, showing the two opposite directions of change. **(D)** Mean strength difference across areas resulted significantly increased in EP3R in patients compared to healthy controls, and significantly decreased in EP3NR patients. *** stands for p<0.001**. (E)** To improve visualization, optimal values of Δ FC strength have been projected on brain surface. It is possible to notice the consistency of the opposite changes, with the majority of areas shifting in the same direction within each condition as compared to controls.

#### Node strength differences

In each condition we assessed the connectivity strength of each area with the rest of the brain (**Fig. 2B** **and Supplementary Fig. S2**). When compared to those of controls, the strength of a few selected areas resulted to be different in patients, following the same trend observed with pairwise FC in groups **(EP3R, EP3NR)** when compared to HC. However, no individual difference was significant after multiple comparison correction. Notably, no discernible trend was found in EP2 patients **(Supplementary Fig. S5A and B).**

To assess whether these trends reflected a globally consistent change in functional strength, we compared the distributions of group average values of strength in each area between conditions. EP3R patients exhibited a significantly increased mean strength compared to healthy controls, indicating higher functional connectivity [*FCS* ^HC^= 39.45±0.14, *FCS* ^EP3R^= 41.97±0.32 (mean ± SE, across areas and subjects), p<0.001]. On the other hand, EP3NR patients, exhibited a strongly significantly decreased mean strength compared to healthy controls, indicating lower functional connectivity [*FCS* ^HC^= 39.45±0.14, *FCS* ^EP3NR^= 37.40±0.35 (mean ± SE, across areas and subjects), p<0.001] (**Fig. 2C**).

For both conditions this change was consistent across areas, with the majority of nodes showing the same direction of change as compared to controls (**Fig. 2E**). Moreover, an overall strength difference was tested for significance by computing the mean functional strength difference across brain areas (Δ*FCS*^EP,HC^) (**Fig. 2D**). EP3R patients exhibited a strongly significantly positive difference [Δ*FCS*^EP3R,HC^= 2.52 ± 0.02 (mean ± SE, across areas), p<0.001, Cohen’s d=1. 3]. On the other hand, EP3NR patients, exhibited a strongly significantly negative difference, indicating lower functional connectivity [Δ*FCS*^EP3NR,HC^= -2.05 ± 0.02 (mean ± SE, across areas), p <0.001, Cohen’s d=0.75].

### 3.3. Model healthy controls

The second step in our analysis consisted of fitting a whole-brain computational model to the healthy resting-state BOLD dynamics to investigate the interplay between structural and functional properties on the healthy brain network. We therefore looked for the optimal combinations of parameters (global coupling G and bifurcation parameters *α*_1_, …, *α*_*N*_) that, given the network’s structure *C* = (*C*_*ij*_), could best reproduce the functional dynamics observed in the empirical data. In **Fig. 3A**, we can see how the optimal combination of parameters allowed to reproduce pretty accurately the empirical functional dynamics of the healthy condition, as indicated by a minimal KS distance of KS = 0.3 ± 0.1 (median ± interquartile range, across 300 simulations). In particular, the optimal value of global coupling was found to be G = 1.38± 0.07 (mean ± SEM, across 300 simulations).

**Fig.3:**
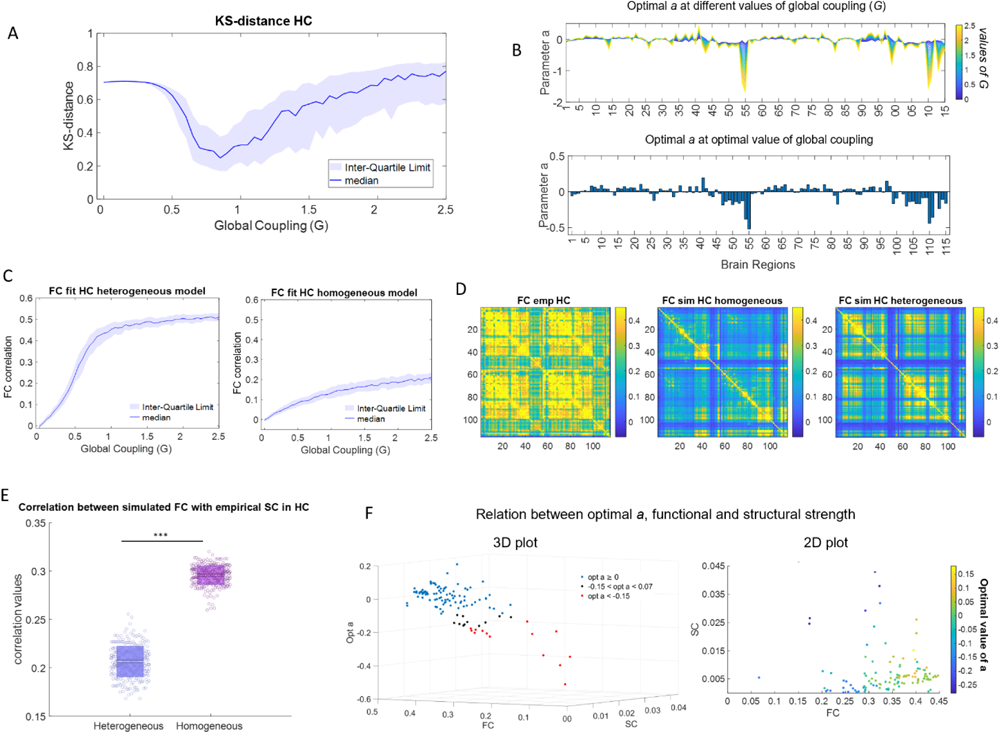
Whole-brain model of HC: Structural and functional information from the empirical data were used to build a whole-brain dynamical model optimised to reproduce HC functional connectivity properties by adjusting one global parameter (the global coupling G) and N local parameters (the bifurcation parameters α_*i*_^*HC*^, …, α_N_), one per node). **(A**) Fitting curve of the model as a function of G, measured with the Kolmogorov-Smirnov (KS) distance between the empirical and the simulated FCD distributions. For each value of G, we used the bifurcation parameters α_1_, …, α_N_, which optimized the power spectral density (PSD) of the simulated BOLD signals in the range 0.04-0.07 Hz to that of the empirical signals. The solid line and the shaded area represent the median and the inter-quartile range, respectively, across N=300 simulations. The optimal combination of parameters (optimal G with its associated optimal α = (α_1_, …, α_N_)) was the one that minimised the mean KS distance between the empirical and the model FCD distributions. **(B)** Optimal value of a for each brain area as the global coupling G varies from 0 to 2.5 (top panel) and at the optimal value of G (bottom panel). **(C)** Fitting curve of the model as a function of G, measured with the Pearson correlation between the empirical and simulated functional connectivity (FC) matrices with the heterogeneous (left panel) and the homogeneous models. In the heterogeneous model, bifurcation parameters α_1_, …, α_N_ were optimized based on the power spectral density of the BOLD signals. In the homogeneous model, all bifurcation parameters were set to α_1_ = ⋯ = α_N_ = 0. The solid line and the shaded area represent the median and the inter-quartile range, respectively, across N=300 simulations. **(D)** Group averaged FC matrices comparing empirical data (left panel), simulated data generated from the heterogeneous (central panel) or homogeneous (right panel) model at the respective optimal value of G. **(E)** Correlation between simulated FC matrix and empirical SC of healthy controls in the homogeneous and heterogeneous model. We can see how the correlation is significantly decreased wen heterogeneity is included in the model, showing how it allows to escape structural constraints in the generation of functional pattern. **(F)** Relation between the optimal bifurcation parameter α ^HC^_j_ (average across 300 simulations), the mean functional strength FCS^HC^_i_and the mean tructural strength FCS^HC^_i_ of each node i (average across subjects). Each dot represents one of the 115 areas. In the right panel the 3 variables are represented in a 3D plot with α ^HC^_i_ on the x axis, FCS^HC^_i_ on the y axis and ^HC^_i_FCS^HC^_i_ on the z axis. To improve visualization, areas associated to a negative value between one and two median absolute deviation from the median (-0.07<α ^HC^_i_<-0.15) are plotted in black, while areas associated to a negative value more than two median absolute deviations from the median are plotted in red (α ^HC^_i_< - 0.15). In the left panel the relation is illustrated in a 2D plot with FCS^HC^_i_ and FCS^HC^_i_ variables on the x and y axes, respectively, ^HC^_i_ and values of optimal a represented as a gradient of colors. Blue indicates negative values and yellow positive values.

The optimal value of (*α*_1_, …, *α*_*N*_) is expressed in a vector represented in **Fig. 3B**. It is interesting to notice that the optimal values of *a_i_* are not homogeneously distributed across different ROIs. In particular, while most regions are associated with optimal *a_i_* parameters around the bifurcation point (*a_i_*= 0), a subset of ROIs showed instead subcritical (*i.e.*<0) values of optimal *a_i_*. These results are in line with a previous work, where the model was applied to a different dataset of healthy controls resting state, indicating the robustness and generatability of the findings ^37^. The median optimal bifurcation parameter for the optimal value of global coupling was found to be *α* = 0.00 ± 0.08 = (median ± MAD, MAD= median absolute deviation, across N=115 areas). To define the subset of areas associated with strong negative values of bifurcation and investigate relevant connectivity properties, we defined a threshold and selected areas associated with a bifurcation parameter more than two MADs away from the median (**Supplementary Table 3**).

#### 3.3.1 Local-global structural-functional interplay and the role of heterogeneity

To explore the interaction between global and local network properties, we examined the relationship between the global coupling parameter G and the local bifurcation parameters *a_i_*. In the top panel of **Fig. 3B**, it’s evident that -along the range of G values that we explored-optimal values of a tend to scale proportionally with the associated value of global coupling. This suggests a compensatory mechanism for states of over or under connectivity. In particular, to higher coupling values correspond greater heterogeneity in the values of parameter *α* across nodes, with values progressively further from the bifurcation point.

To assess the role of heterogeneous negative values of bifurcation parameters, we attempted to model healthy brain functional dynamics by imposing a homogeneous value of *a_i_*= 0 (edge of criticality) in all regions of interest (ROIs) and compared it to the previous model with heterogeneous parameters. The model with homogeneous parameters resulted in significantly worse FC fit. It failed to reproduce essential spatial connectivity properties of the functional network, as indicated by a significantly lower Pearson correlation between empirical and simulated FC compared to the heterogeneous model (**Fig. 3C**). Specifically, the homogeneous model struggled to replicate the hierarchical organization of the network into communities (**Fig. 3D**). Moreover, when homogeneous bifurcation parameter values were imposed, the correlation between optimal simulated FC and the underlying empirical structural connectivity (SC) significantly increased compared to when heterogeneity was allowed (**Fig. 3E**). This, along with earlier observations, suggests that local heterogeneities may enable the network to sustain more complex spatial patterns of FC while (1) avoiding constraints imposed by SC and (2) preventing states of under or over synchronization as structural connectivity strengthens^38^.

Furthermore, using a generalized linear model, we explored the relation between optimal values of parameter *a_i_* in each area and their corresponding structural and functional strength. The combination of *FCS*_*i*_^HC^, *SCS*_*i*_^HC^and the interactions between them, explained 64.2% of the variance in optimal *a_i_*, values, as shown in **Fig. 3F**. Specifically, the optimal bifurcation parameter value was positively correlated with functional strength (r=1.06, p<0.01), which explained 44.0% of its variance across regions, and negatively correlated with structural strength (r=-5.83, p<0.01), which explained 15.7% of its variance across regions. In particular, strong negative values can be associated with either 1) areas characterised by low functional strength and low structural strength, indicating disconnection from the rest of the network, or 2) areas with very high structural strength, often referred to as connectivity hubs (**Fig. 3F** **right panel**).

### 3.4 The role of local negative bifurcation parameter in a network

To better understand the effect of the local bifurcation parameter on network dynamics and how this effect is modulated by the structural strength of the node, we built a simplified model with realistic parameters and tested a number of different scenarios. We hypothesised that a negative value of the bifurcation would decrease node’s sensitivity to incoming stimuli, thereby altering signal propagation and reducing global synchronization. To assess our hypothesis, we created a network consisting of two interconnected communities linked through a central hub. We manipulated the bifurcation parameter associated to this hub and applied gradually increasing synchronization stimuli to it.

We found that as the hub’s *a* value became more negative (subcritical), it required stronger perturbations to achieve the same global synchronization observed at *a*=0 (**Fig. 4A**).

**Fig.4:**
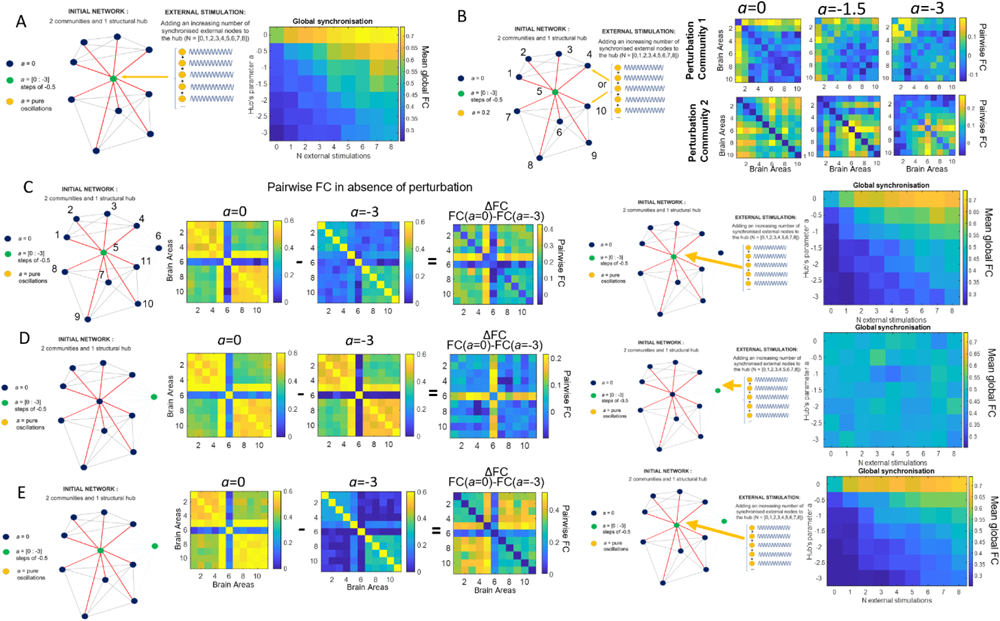
Toy model: Simplified network of coupled Hopf oscillators. **(A, B)**. Effect of a negative value of a associated to a structural hub. The left panel shows a network organised into two communities connected through a single node (hub). The strength of structural connection of each node toward the other nodes within the same community was set to 0.12 (grey lines), while that of the hub with other nodes was set to 0.2 with all nodes in the network (red lines). The bifurcation parameter was initially set to be equal to *α* = 0 in all nodes. Subsequently, the bifurcation parameter assigned to the hub was modified to be increasingly negative (green dot). **(A)** Effect of an external perturbation on the hub. For each value of bifurcation parameter, we connected to the hub an increasing number (1 to 8) of synchronised external nodes. We then computed the global functional connectivity at each iteration (right panel). **(B)** Effect of an external perturbation on one network community. For each value of bifurcation parameter, we connected to a node either in the first (top right panels) or second (bottom right panels) community an increasing number of synchronised external nodes. We then computed the pairwise functional connectivity at each iteration and plot the connectivity pattern resulting at maximum perturbation. **(C, D, E)**. Effect of a negative value of a associated to very disconnected nodes. We included an additional node weakly connected to the original network, whose strength of structural connectivity with the rest of the network was set to 0.01. The bifurcation parameter was initially set to *α* = 0 in all nodes (left panel). Subsequently, we evaluated tree different scenarios: the bifurcation parameter assigned to the hub **(C),** to the disconnected node **(D),** or to both **(E)** was modified to be increasingly negative (green dot). For each scenario, we then computed pairwise functional connectivity at each iteration in absence of perturbation (spontaneous dynamics). To highlight the heterogeneous influence of the bifurcation parameter on network dynamics, we additionally plot the difference in pairwise connectivity between the condition were *α* was set to extremely negative values as compared to *α* = 0 (central panels). Finally, for each value of the bifurcation parameter, we connected to the hub **(C, E)** or to the disconnected node **(D)** an increasing number of external synchronised nodes (perturbed dynamics), and we computed the global functional connectivity at each iteration (right panel).

We then applied the perturbation to a node within a community and observed its spread through the network. At *a*=0, the synchronization wave spread throughout the entire network. However, with more negative *a* values, the synchronization wave remained more confined to the perturbed community, affecting the rest of the network less (**Fig. 4B**).

We also examined the effect of a negative *a* value on very weakly connected nodes. We added such a node to the network, (**Fig. 4C-E**), and tested three scenarios:

1. We assigned progressively more negative values of *a* to the hub. As a result, the hub progressively disconnected from the network, and leaded to a homogeneous reduction in pairwise functional connectivity among the other nodes. The disconnected node remained unaffected (**Fig. 4C**). To compare it to the previous case, we also added a progressively increasing number of synchronized perturbations to the hub and computed the mean FC of the network, obtaining similar results as before.
2. We assigned progressively more negative values of *a* to the disconnected node. This caused the node to progressively disconnect from the rest of the network. However, this had minimal impact on the rest of the network, and even increasing the perturbations applied to the disconnected node, its influence remained limited.
3. We assigned progressively more negative values of *a* to both the hub and the disconnected node. In this case, the network’s response to synchronization perturbations applied to the hub was similar to scenario 1. However, in the spontaneous regime, the pairwise dynamics were significantly altered by the additional manipulation of *a* in the disconnected node. In particular, connections between communities were more reduced than connections within each community, resulting in a more defined segregation of the network compared to scenario 1 (**Fig. 4E**).

Overall, these findings suggest that negative bifurcation parameters in key network nodes, whether highly connected or disconnected, play a crucial role in a) filtering out irrelevant stimuli and preventing excessive network synchronization, b) regulating the flow of stimulus waves within and between structured communities, and c) facilitating the emergence of intricate patterns of functional connectivity with an appropriate degree of segregation.

### 3.5 Model EP patients

To gain insights into the possible mechanisms underlying the empirical changes in functional connectivity reported in the previous sections, we built independent whole brain models to reproduce the spatiotemporal dynamics observed in the resting brain activity of the two subgroups of EP subjects (EP3R, EP3NR). We observed a significant reduction in the optimal global coupling parameter (G) in patients from stage III, both remitting-relapsing and non-remitting, compared to controls (HC optimal G = 1.38 ± 0.07; mean ± SEM across 300 simulations), as indicated by the shift in the KS distance curve (**Fig. 5A**).

**Fig.5:**
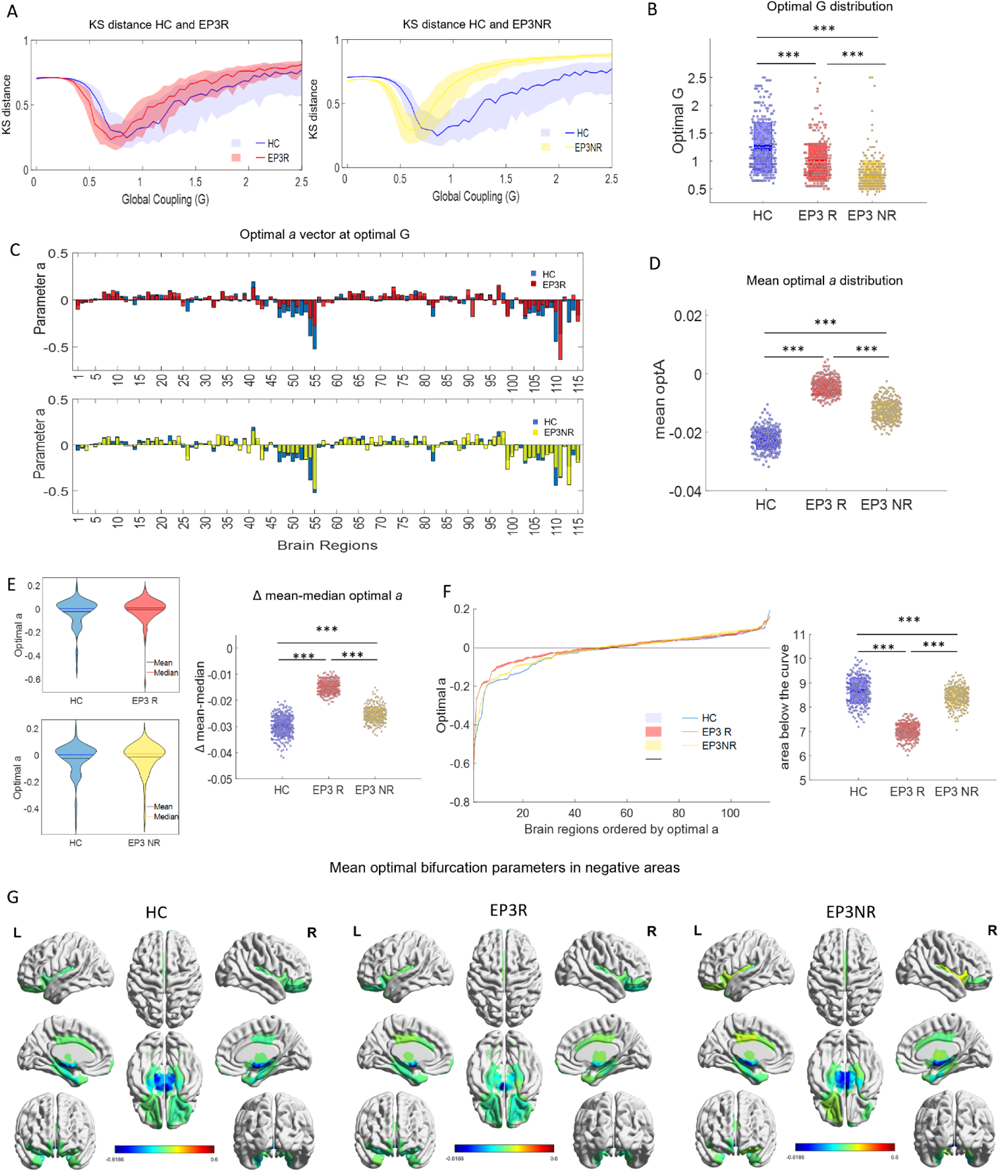
Whole-brain model of patient groups EP3R and EP3NR compared to HC: *Structural and functional information from the empirical data were used to build two whole-brain dynamical models optimised to reproduce functional connectivity properties of EP3R and EP3NR, respectively, by adjusting the global parameter G and the bifurcation parameters. In all panels, HCs are represented in blue, EP3R in red, and EP3NR in yellow. Differences between groups were assessed using Wilcoxon rank-sum test followed by Bonferroni p-value correction (p<0.001). **(A)** Fitting curve of the model as a function of G, measured with the Kolmogorov-Smirnov (KS) distance between the empirical and the simulated FCD distributions. The solid line and the shaded area represent the median and the inter-quartile range, respectively, across N=300 simulations. The optimal combination of parameters (optimal G with its associated optimal α* = (*α*_1_, …, *α*_*N*_)*) was the one that minimised the mean KS distance between the empirical and the model FCD distributions. **(B)** Optimal global coupling G distribution. Each dot represents a simulation, and boxplots represent the mean of the measures’ values with a 95% confidence interval of the mean (dark) and 1 SD (light). **(C)** Mean optimal value of a* for each of the 115 nodes *at the optimal value of G in HC and in each EP group (average across 300 simulations). **(D)** Distribution of mean optimal bifurcation parameters across brain regions at optimal global coupling G. Each dot represents a simulation. Boxplots represent the mean of the measures’ values with a 95% confidence interval of the mean (dark) and 1 SD (light). **(E)** Comparison of mean and median values of optimal bifurcation across brain regions between conditions. Violin plots on the left show optimal bifurcation parameters distribution across brain regions. Mean is plotted as a black line, while median is plotted as solid coloured line. Boxplots on the right represent the difference between mean and median in each condition. Each dot represents a simulation. Boxes represent the mean of the difference across simulations with a 95% confidence interval of the mean (dark) and 1 SD (light). **(F)** Left panel: optimal value of parameter a across simulations for each of the 115 nodes sorted by node’s optimal a in each condition, at the optimal value of G* (solid and shaded lines represent mean and SEM across N=300 simulations)*. Right panel: distribution of areas under the curve of optimal a in each condition, illustrating loss of heterogeneity and the shrink toward values around zero in patients. *** stands for p<0.001. **(G)** To improve visualization, optimal values of bifurcation parameters in ROI’s originally associated with negative have been projected on brain surface. It is possible to notice a decrease of highly negative values (dark blue) in patients, especially in EP3R, as compared to controls*.

In particular, the optimal G was 1.06 ± 0.05 for the EP3R subgroup and 0.72 ± 0.03 for the EP3NR subgroup, with a significant difference from controls (unpaired t-test, p<0.001). Notably, the reduction of optimal global coupling, was significantly larger in magnitude in EP3NR than in EP3R (unpaired t-test, p<0.001) (**Fig. 5B**).

We then compared the optimal bifurcation parameters across nodes within and between conditions and observed a complex pattern of alterations (**Fig. 5C**). EP3 patients displayed less heterogeneity across areas and tended to have bifurcation parameters closer to criticality (*a* = 0) compared to HCs. This, according to results presented in previous sections, implies that ROIs’ activity in EP3 patients is less stable and more sensible to incoming stimuli than in HC. It also implies increased constraints imposed by structural connectivity on the emergence of functional patterns. This was especially the case for the dynamics of ROIs associated with strong negative values of *a* in HC, whose role in regulating network dynamics was established in section 3.4. To quantify this change, we computed the mean of optimal bifurcation parameters across brain regions and compared it between conditions, observing a significant shift toward less negative values associated with a reduction in heterogeneity across areas in EP3R [-0.0126 ± 2e-4] (grand mean ± SE across repetitions, p<0.001) and EP3NR [-0.0126 ± 2e-4] (grand mean ± SE across repetitions, p<0.001) as compared to healthy controls [-0.0225 ± 2e-4] (grand mean ± SE across repetitions). This alteration was more extensive in EP3R than in EP3NR (p<0.001) (**Fig. 5D**).

Since the change affected mainly ROIs originally associated with extremely negative values, we compared the mean and median values of bifurcation parameters across brain regions in each condition, knowing that the median is less influenced by extreme values. As expected, in HC we observed a consistent difference between the two measures, with the mean being considerably shifted toward negative values [-0.030 ± 0.004] (mean Δ ± STD across repetitions). In EP3R, where negative values were less extreme, the difference between these two statistics was significantly reduced [-0.015 ± 0.003] (mean Δ ± STD across repetitions p<0.001). The same alteration could be seen in EP3NR, [-0.025 ± 0.002] (mean Δ ± STD across repetitions p<0.001), but it was significantly less consistent than in EP3R (p<0.001), as can be observed in **Fig. 5E**.

The loss of heterogeneity and the shrink towards criticality (*a*=0) can also be observed by the reduction of the area under the curve of the optimal bifurcation parameter across brain regions illustrated in **Fig. 5F**. This area was in fact significantly smaller in both EP3R [7.02 ± 0.02] (mean ± SEM across repetitions, p<0.001) and EP3NR [8.44 ± 0.02] (mean ± SEM across repetitions, p<0.001), as compared to HC [8,67 ± 0.03] (mean ± SEM across repetitions). Consistent with our previous findings, the reduction was significantly more consistent in EP3R than in EP3NR (p<0.001). To improve visualization, for each condition optimal values of bifurcation parameters of ROI’s originally associated with negative values have been projected on brain surface (**Fig. 5G**). Furthermore, in line with previously discussed results indicating how a loss in heterogeneity corresponds to increased SC constraints imposed on the emerging FC patterns (**Fig. 3E**), we observed that in EP3R patients’ empirical FC and SC tend to correlate more as compared to HC, even though not sufficiently to reach significance **Supplementary Fig. S7**. Additionally, in **Supplementary Fig. S3,** we show how the change in the optimal value of bifurcation parameter in each node is related to its functional and structural strength. As a further step we investigated whether the change in the optimal value of the bifurcation parameter across areas (Δ*α*^EP,HC^) correlated with the change in functional strength (Δ*FCS*^EP,HC^) reported in section 3.2 and illustrated in **Fig. 2**. In EP3R, but not in the other two subgroups, we found a significant correlation between these two metrics (r=0.33; p=0.02) (**Supplementary Fig. S4**).

Finally, in **Supplementary Fig. S5,** we show that empirical differences in FC measures between conditions were replicated in simulated data, validating the model’s ability to capture relevant properties of the data.

### 3.6. Additional analysis on EP2

Despite not including patients from stage II in the main analysis due to heterogeneity of the group that would complicate interpretation of the results, we repeated the whole pipeline of analysis with this subgroup in order to assess whether the findings observed in EP3R and EP3NR could be generalised to all early psychotic patients. None of the alterations found in any of the two EP3 subgroups could be detected in EP2, proving once more the importance of clinical staging to address heterogeneity within psychotic patients.

## 4. Discussion (1718 to 1350)

In this work we propose to classify patients according to their ability to remit from a first psychotic episode, aiming to investigate the neural correlates underlying these different clinical profiles and outcomes (**Fig. 1A**). In fact, while neither of these patients have fully recovered, stage III psychosis patients present different clinical pictures at the moment of the scan (i.e., temporary reduction of symptoms for the remitting-relapsing patients or residual symptoms for the non-remitting ones). We therefore analysed fMRI data of remitting-relapsing and non-remitting stage III patients and compared their respective functional connectivity profiles with those of healthy controls (**Fig. 1B**), highlighting two opposite trends of alteration, which suggests the existence of a compensatory mechanism. A whole brain model (**Fig. 1C**) allowed us to investigate hidden correlates underlying these alterations, highlighting some crucial properties of the healthy network and how those are affected in the two subgroups of patients.

### 4.1. Opposite functional connectivity impairments in remitting-relapsing and non-remitting patients

We hypothesized that different functional neural patterns underlie distinct clinical profiles in two subgroups of stage III patients. Consistent with our expectations, significant differences in functional strength were observed, with opposite alterations were found when comparing brain activity in these two subgroups with that of healthy controls. Patients with residual symptoms after the FEP, who had therefore not completely remitted at the moment of the scan (EP3NR), exhibited a significant reduction of mean functional strength across areas (**Fig. 2C, D**), aligning with previous findings of reduced structural strength ^30^. In contrast, patients from stage III that were able to remit after their psychotic episode, despite potentially having subsequent relapses (EP3R), showed increased mean functional strength, indicating a potential compensatory mechanism (**Fig. 2C, D**). These opposing patterns of alterations were not only observed in mean values of functional strength across brain areas, where statistical significance was reached, but were also evident as a trend in pairwise connectivity (**Fig. 2A**) and individual area strength (**Fig. 2B**). The majority of areas showed consistent shifts within each patient subgroup and these shifts were in opposite directions between the two subgroups as compared to controls (**Fig. 2E**).

The significant differences observed between remitting-relapsing and non-remitting patients, often analysed as a combined group, could contribute to the heterogeneity in functional connectivity studies of psychotic disorders ^23^. We therefore suggest that further investigations on this topic should take into account this source of intra-patient variability. Moreover, our findings suggest the presence of a potential compensatory mechanism in patients who achieve remission. The increased functional connectivity observed, previously reported in psychotic disorders ^26,39,40^, especially in early stages, may represent an over compensatory attempt to restore clinical functioning by rebalancing the structural-functional interplay ^41,42^. As discussed by Fornito and Bullmore, the dissociation between reduced structural connectivity and increased functional connectivity can be interpreted as a neural activity differentiation, involving a disruption of the typical segregation of neural functions or as a compensatory process. It is also possible that both mechanisms contribute to this phenomenon ^23^.

### 4.2. Properties of structural-functional interplay in the healthy brain

We investigated our hypothesis exploring the interplay between structural, dynamical, local, and network properties of the healthy brain with a computational whole brain model (**Fig. 1C**, **Fig. 3**). This model incorporates empirical structural information obtained from DSI measures and replicates the functional dynamics observed in healthy individuals by adjusting two key factors: (i) the global coupling parameter (G), controlling overall effective strength of connections, and (ii) regional bifurcation parameters (*a*), describing each brain region’s local behaviour.

Previous research has established that the brain operates optimally in a criticality regime, where it teeters on the edge of a bifurcation point between noisy and oscillatory behaviours ^43^. This regime fosters the emergence of diverse and complex metastable activity patterns, ensuring a balance between stability and flexibility ^44,45^. However, recent studies emphasized the importance of local heterogeneity in node behaviours for maintaining global criticality level^46^. Our study supports this idea by showing that the healthy brain’s dynamics are best represented when specific brain regions are assigned negative values for the bifurcation parameter, indicating locally stable yet noisy behaviour (**Fig. 3B**).

To investigate these nodes’ properties, we applied a threshold to identify areas with bifurcation parameters deviating by more than two absolute deviations from the median and extracted their values of structural and functional strength. Our results reveal a strong correlation between bifurcation parameter values across nodes and combined functional-structural properties (**Fig. 3F**). This suggests that the bifurcation parameter is influenced by both dimensions, hinting at its role in regulating their interplay. Additionally, using a toy model, we shed light on the role of negative bifurcation parameters in enhancing node stability by dampening incoming stimuli, both internal and external. This property is especially relevant in nodes with prominent network characteristics like connectivity hubs and isolated nodes, as it prevents excessive network synchronization while allowing optimal segregation and the emergence of complex functional connectivity patterns (**Fig. 4**). Moreover, heterogeneity in bifurcation parameters can relax structural constraints and promote greater variety and flexibility in functional dynamics ^38^, as evidenced by the increased functional-structural correlation observed when homogeneous values of bifurcation parameters are imposed on the model (**Fig. 3C-E**).

### 4.3. Alterations in network properties and potential compensatory mechanisms in EP patients

Our investigation then delved into how these properties were altered in patients and whether they could account for the observed shifts in functional dynamics. To do this, we independently constructed whole-brain models using the structural connectivity matrices of both remitting-relapsing and non-remitting patients (**Fig. 5**).

Firstly, we found that all stage III patients exhibited significantly lower optimal coupling strength (G) compared to controls (**Fig. 5A,B**), indicating reduced structural strength, typically associated with diminished conductivity effectiveness. This reduction was significantly more pronounced in non-remitting-relapsing patients (EP3NR), suggesting a more severe initial impairment. This alteration might signify a global decrease in long-range excitatory connectivity (E-E), in line with the structural dysconnectivity hypothesis ^19^. Possible contributors to this decrease could include alterations in myelination, global changes in synaptic density or strength, and/or widespread modifications in receptors ^47,48^.

Furthermore, stage III patients displayed a loss of heterogeneity in optimal local bifurcation parameters compared to controls, converging toward values around zero (**Fig. 5C-G**). According to our toy-model study, this implies heightened sensitivity to incoming stimuli and reduced stability. Notably, this alteration was more prominent in remitting-relapsing patients (EP3R). The less pronounced reduction in connectivity coupling in this subgroup, combined with a more extensive alteration of local node properties, may account for the observed over-compensatory increase in functional connectivity among remitting-relapsing patients, as opposed to non-remitting patients. Intriguingly, only in the EP3R group these alterations in bifurcation parameters significantly correlate with changes in empirical functional strength (**Supplementary Fig. S4**), suggesting their potential role in underlying connectivity changes in this condition. Bifurcation parameters can be interpreted as encoding the local responsiveness (or resilience) ^37^ of brain regions to incoming stimuli, as shown with the toy-model study. The loss of heterogeneity observed in patients, particularly in the remitting-relapsing group, could reflect a loss of functional flexibility due to increased structural constraints ^49^, which may have clinical implications, particularly in the long term. In particular, as discussed in previous works, this could contribute to impairments in cognitive flexibility reported in patients^50^.

We hypothesize that these changes may reflect heterogeneous local imbalances between excitatory and inhibitory activity, aligning with the disinhibition hypothesis ^51,52^. Moreover, this aligns with the hypothesis proposed by Krystal et al. in a recent insightful review ^42^. The review suggests that increased intrinsic excitability or reduced inhibitory tone may serve as allostatic adaptive compensatory mechanisms in response to NMDAR related connectivity alterations, potentially leading to functional and structural consequences. Further empirical investigations are needed to test the validity of this hypothesis and provide a deeper understanding of the underlying mechanism. In line with our results, Horne et al. observed differentially disrupted connectivity in treatment-resistant and -responsive patients with schizophrenia. In particular, they found that responsive patients display effective compensatory increased top-down connectivity from ACC to sensory regions serving to reduce sensory input to the striatum (control of sensory precision during the task) and an absence of this compensatory cognitive control mechanism in resistant patients ^53^.

Critically, brain data from patients in stage II, who also experience remission from the first episode similar to EP3R but have not relapsed at the time of the scan, did not exhibit any of the alterations observed in the EP3R group (**Supplementary Fig. S6**). They did not show significant differences compared to healthy controls in terms of empirical measures, and most importantly, they did not demonstrate the changes in the parameters observed in stage III patients. Interestingly, even when some parameter alterations were detected in this group, they consistently displayed an opposite trend in both global coupling and bifurcation parameters. This suggests a disease progression component within this compensatory mechanism. However, the heterogeneity of patients in group II, comprising individuals who have not yet relapsed but may do so in the future, as well as those who will maintain full recovery, complicates further interpretation.

Taken together, these findings suggest that imbalances in the interplay between structural and functional components may contribute to the underlying pathophysiological processes of early psychosis, providing insights into their clinical implications. Additionally, these findings highlight how heterogeneous alterations in local node properties can manifest as global network changes, emphasizing the significance of maintaining local and global equilibria. Importantly, the observed heterogeneity in local alterations may also arise from the preferential vulnerability of specific nodes or networks to global alterations, such as changes in synaptic NMDAR function, as suggested by Yang et al. ^54^

### 4.4. Limitations and future perspectives

One limitation of this study is the small sample size, especially when dividing patients into subgroups, although it is comparable to similar studies. To ensure the generalizability of the findings, replication on larger datasets is necessary. Future research should adopt patient-level analysis and simulations for valuable clinical predictions, and incorporate biologically-constrained models with finer details of network properties, for a more accurate interpretation of results. Mesoscopic or microscopic models of connectivity and synaptic plasticity can provide deeper insights into underlying processes driving the compensatory mechanism ^55–58^. Furthermore, longitudinal approaches will be essential to comprehend alterations’ long-term evolution, disease progression, and prognosis, and results validation through machine learning predictions can offer clinical relevance insights for personalized care. Finally, consideration of medication’s influence on brain dynamics is crucial; however, our analysis primarily focuses on differences between remitting-relapsing and non-remitting patients, with comparable medication levels to mitigate confounding effects.

## Funding

The project that gave rise to these results received the support of a fellowship from “la Caixa” foundation “(ID 100010434)”. The fellowship code is: “(LCF/BQ/DI19/11730048)”, and financed L.M work. In addition, G.D., M.V.V. and L.M. were supported by the Human Brain Project Specific Grant Agreement 3 Grant agreement no. “(945539)” financed by the European Commission and by the Spanish Research Project ref. “(PID2019-105772GB-I00/AEI/10.13039/501100011033)”, financed by the Spanish Ministry of Science, Innovation and Universities (MCIU), State Research Agency (AEI). This last institution also financed M.V.V trough the Grant/Award Number: PID2020-119072RA-I00/AEI/10.13039/501100011033. A.L.G. was supported by Swiss National Science Foundation Sinergia grant no. 170873. L.A.E. was supported by a research grant of University of Lausanne, Switzerland. L.A. was supported by a fellowship of the Adrian and Simone Frutiger Foundation and from Carigest Foundation. P.K. was supported by a fellowship from the Adrian and Simone Frutiger Foundation. Y.A.G. and P.H. were financially supported by Swiss National Science Foundation grant #320030-197787.

## AUTHOR CONTRIBUTIONS

R.J., P.B., P.K., P.C., L.A., and L.A.E. collected data and performed clinical evaluation; PB and PK assessed the clinical staging; P.H, and Y.A.G. collected and pre-processed the neuroimaging data; A.L.G., L.M, M.V.V. and G.D. designed the research; L.M., A.L.G. and M.V.V. analysed the data; L.M. and M.V.V. wrote the manuscript; M.V.V and G.D. supervised the research. All authors contributed to the editing of the manuscript. Correspondent author: L.M.

## Competing interests

‘The authors report no competing interests.’

## Notes

### Competing Interest Statement

The authors have declared no competing interest.

### Summary of Updates

author affiliations updated; author contributions updated

