## Supplementary data for "Brain network connectivity underlying remission in early psychosis: a whole brain model approach"

### SUPPLEMENTARY METHODS

#### 1. Participants

For the current work we included a total of 216 subjects (88 early-psychosis patients, EP, and 128 healthy controls, HC), in the range from 18 to 35 years *[26.3 ± 6 (mean ± SEM)*]. All participants provided informed written consent for this study, and the procedure was approved by the local ethics Committee.

The 88 EP patients (26.0 ± 6.2 yo) were recruited from the Treatment and Early Intervention in Psychosis (Tipp) program, an early intervention program of the Lausanne University Hospital, Switzerland which offers 3 y of treatment to patients aged 18–35 years. Inclusion criteria for the TIPP programme were (i) meeting the psychosis threshold as defined by the Comprehensive Assessment of At-Risk Mental States, (ii) no antipsychotic medication for >6 months, (iii) no psychosis related to intoxication or organic brain disease, and (iv) intelligence quotient ≥70 [60].

A total of 128 age-, gender- and handedness-matched HC were recruited from the same catchment area. HC were not affected by mood, psychotic or substance use disorder and had no first degree relative with a psychotic disorder [61].

Exclusion criteria for both patients and controls were history of neurological disorder, severe head trauma or mental disability (IQ<70).

#### 2. Diagnosis, clinical variables, and staging

The patients were initially stratified into four distinct groups (stages II and IIIa-c (**Fig. 1**)) based on a consensus assessment by two experienced psychiatrists, according to the clinical staging model proposed by McGorry et al. 2006. Subjects in stage II (EP2) were first-episode psychosis patients, with one psychotic episode according to the CAARMS psychosis-threshold subscale and no past episodes at the time of the study (i.e., discrete disorder).

Patients in stage III were defined as follows:

- IIIa: incomplete remission from stage II at 12 months after entry to care and following a reasonable course of treatment (>3 months), with duration of illness no longer than 5 years;
- IIIb: relapse of a psychotic episode (i.e., discrete disorder has fully remitted but then relapsed to the full extent described in stage II);

- IIIc: two or more relapses after stage II with remission between episodes.

Because the goal of the study was to investigate neural mechanisms critically involved in remission from the acute FEP and because we considered that the elevated heterogeneity of stage II patients would complicate interpretation of the results, we restricted the main focus of the current study to the analysis of stage III patients. In particular, for the current work, patients from stage IIIb and IIIc were grouped together into a subgroup of remitting-relapsing patients (EP3R), including 31 patients in total, while patients from stage IIIa were kept separately in a subgroup of non-remitting patients (EP3NR), including 20 patients.

#### 3. Imaging

##### *MRI acquisition*

MRI data was acquired using two different 3-Tesla scanners, namely the Magnetom TrioTim and PRISMA from Siemens Medical Solutions. Each scanner was equipped with a 32-channel head coil. The MRI sessions consisted of three sequences: a magnetization prepared rapid acquisition gradient echo (MPRAGE) T1-weighted sequence, a spin-echo echo-planar imaging (SE-EPI) diffusion spectrum imaging (DSI) sequence and a gradient echo-planar imaging (EPI) sequence sensitive to BOLD (blood-oxygen-level-dependent) contrast (resting-state functional magnetic resonance imaging). The MPRAGE-T1w images were obtained with the following parameters: echo time (TE) = 2.98 ms, repetition time (TR) = 2300 ms, inversion time (TI) = 900 ms, flip angle (FA) = 8°, field of view (FOV) = 160 x 240 x 256 mm$^3$, and voxel size = 1 x 1 x 1.2 mm$^3$.

The DSI sequence (using the q4half acquisition scheme) included 1 b0 acquisition and 128 diffusion weighted directions. The DSI parameters were as follows: TE = 103 ms, TR = 5900 ms, FOV = 211 x 211 x 114 mm$^3$, voxel size = 2.2 x 2.2 x 3 mm$^3$, and maximum b-value = 8000 s/mm$^2$.

In this study, resting-state functional magnetic resonance imaging (rs-fMRI) data was acquired using a standardized scanning sequence. The EPI sequence employed a TR= 1920 ms and TE = 30 ms, optimizing the sensitivity to blood-oxygen-level dependent (BOLD) contrast, which reflects changes in neural activity. Thers-fMRI acquisition had an isotropic 3.3mm voxel size, with a 0.3mm inter-slice gap and covering a total of 64 × 58 × 32 voxels. The acquisition times for the MPRAGE-T1w , DSI and rs-fMRI sequences were approximately 7, 13 and 9 minutes, respectively.

Image quality assessment was conducted to ensure the high quality of the dataset. All images included in this study underwent visual inspection by experts from various fields. Our quality control protocol incorporated exclusion criteria that considered incidental findings and poor image quality.

Furthermore, we calculated different quality control metrics using MRIQC and QUAD (QUality Assessment for DMRI), which took into account signal, noise, image smoothness, contrast, and specific artifacts[62–64]. More details about quality control and data harmonization can be found in Alemán-Gómez et al 2022.

##### *Image processing*

In the gray matter parcellation process, individual T1-weighted images underwent skull-stripping using the Computational Anatomy Toolbox (CAT12) and manual correction. The FreeSurfer stream was then utilized for cortical surface reconstruction and subcortical parcellation. The hippocampal subfields and brainstem were segmented using methods proposed by Iglesias et al, while an atlas-based segmentation approach with the Advanced Normalization Tools (ANTS) was employed for thalamic nuclei segmentation. These segmentations were combined with the FreeSurfer outputs to obtain comprehensive gray matter parcellations consisting of 115 regions of interest.

For DSI processing and fiber tracking, individual DSI images underwent a series of correction steps using Mrtrix3 and FSL. These steps included denoising, bias correction, motion correction, eddy current correction, and distortion correction. Diffusion tensors and orientation distribution functions (ODFs) were derived from the corrected DWIs, with ODFs providing higher angular resolution. Deterministic streamline tractography was performed using the sdstream algorithm to map the structural connectivity of the brain.

Resting-state fMRI preprocessing was conducted using the fmriprep pipeline. This involved motion correction, co-registration with anatomical images, spatial normalization, slice-timing correction, smoothing, and nuisance signal regression to remove confounding factors. The preprocessing ensured high-quality data for subsequent functional connectivity analysis.

#### 4. Measures of structural connectivity

The structural connectivity between each pair of cortical and subcortical regions was quantified as the density of streamlines connecting the two regions, which resulted in 115-nodes, weighted undirected brain-networks. Density connectomes take into account not only the presence of streamlines connecting brain regions but also the size of the regions themselves by normalizing the number of streamlines by the geometric mean of the volumes of those regions. This normalization factor accounts for the differences in region sizes, ensuring that larger regions do not have an inherent advantage in terms of connectivity measures. By incorporating this size correction, density

connectomes offer a more accurate representation of the true connectivity patterns between brain regions, unbiased by differences in their sizes.
To compute density connectomes, the number of streamlines connecting each pair of regions was divided by the geometric mean of their volumes, yielding a density measure that reflects the relative strength of connectivity between the regions while accounting for their sizes. This information was saved as a connectivity matrix, where each element represents the density of connections between specific pairs of brain regions. For consistency reasons and to limit possible biases in the network analyses, connections that were present in less than 50% of the subjects (including both healthy controls and patients) were discarded.

#### 5. Measures of functional connectivity

BOLD signal was filtered (0.04-0.07 Hz) in order to focus on the most relevant frequency bands, as demonstrated in previous works [36]. We then computed Functional connectivity (FC) analyses to investigate undirected levels of coupling between pairs of areas across time. Functional connectivity can be computed trough different analysis. For the main analysis described in this work we computed FC as the Pearson correlation between the BOLD signal of each pair of brain areas over the entire recording period. Therefore, for each subject a NxN FC matrix was obtained, where N=115 is the number of brain areas, and the temporal dimension is collapsed into a single value.
To assess the robustness of the results, we repeated all analyses using an independent FC measure which relies on the consistency of phase relationships across time. In this case, BOLD signals are Hilbert transformed to compute the phase of each brain area at each time point (T=276 time points). Then, instantaneous phase coherence (iFC) between a pair of brain areas i and j is obtained by calculating the cosine of the phase difference at any time point t:

$$iFC(i,j,t) = \cos(\theta_i(t) - \theta_j(t)), \quad [1]$$

where $\theta_i(t)$ is the instantaneous phase of the filtered BOLD signal at node i. Note the iFC ranges from -1 to 1. Two brain areas that are in phase (i.e., with no phase delay) have iFC = 1, while signals in anti-phase (phase delay of $\pi$) result in iFC = -1. In contrast, orthogonal signals (i.e., with a phase delay of $\pi/2$) will have iFC = 0. Such analysis resulted in a NxNxT iFC matrix (N=115 areas, T=276 time points) for each subject and each condition. This matrix was subsequently averaged across time to investigate the overall connectivity level of each pair of brain areas, obtaining a NxN FC matrix. Results obtained when computing FC through phase analysis are shown in supplementary results.

#### *Global network-level analyses*

Global functional connectivity measures the overall level of coupling of all areas within the network across time and was calculated as the mean value of the FC matrix. Moreover, we investigated other global properties of the network such as the root-mean-square (RMS) signal amplitude, the segregation and the integration, computed as described in previous works (see supplementary materials and in Deco et al. 2015, 2018; Deco and Kringelbach 2017).

Wilcoxon ranksum test was used to investigate differences in global measures between conditions, and Bonferroni was applied to correct for multiple comparison (3 comparisons across conditions). A threshold of $\alpha = 0.05$ was used to define statistical significance.

#### *Pairwise connectivity and node strength analysis*

In each subject, node strength ($FCs_i$) was calculated as the sum over columns of the pairwise FC matrix and represents the overall level of connectivity of each individual node (i). Group averages were obtained for each pairwise connectivity and node strength by computing the corresponding mean value across subjects in each group. We then investigated differences in pairwise connectivity and in node strength between each EP group (EP2, EP3-NR, EP3-R) and HCs. To test for statistical significance, we used a Wilcoxon ranksum test for each comparison. Bonferroni was applied to correct for multiple comparison in each EP group ($\mathrm{N}^*(\mathrm{N}-1)/2$ comparisons in pairwise connectivity, N comparisons in node strength, with N=115. Threshold of $\alpha = 0.05$ was used to define statistical significance.

Next, we assessed whether there was a consistent change in functional strength across all nodes between each EP group and HCs. For each condition and node i, we used the previously computed group average node strength ($FCS_i^{condition}$). We therefore compared the distributions of strength values across nodes in the different conditions and applied a paired t-test to test for significance**.**

Then, we computed the mean difference of strength in each area. More specifically, for each area i we subtracted the node strength level in HC ($FCS_i^{\mathrm{HC}}$) to the corresponding node strength level in each pathological condition ($FCS_i^{\mathrm{EP3R}}, FCS_i^{\mathrm{EP3NR}}$), resulting in a N-dimensional vector of differences for each EP group:

$$\Delta FCS_i^{\mathrm{EP,HC}} = FCS_i^{\mathrm{EP}} - FCS_i^{\mathrm{HC}} \qquad [2]$$

Finally, we tested whether the obtained global difference of strength between conditions was large enough to reject the null hypothesis that the two conditions had the same mean value (H0: $\Delta=0$). Bonferroni was applied to correct for multiple comparison (condition=3 comparisons) and threshold of $\alpha = 0.05$ was used to define statistical significance. Effect size was assessed with the Cohen's *d*, i.e., by the difference between the two means (each EP group vs HC) divided by the standard deviation of the distribution of differences [68].

##### *Relation between structural and functional connectivity*

We investigated whether the level of functional connectivity associated to each pair of brain areas correlated with the corresponding values of structural connectivity. To this aim, for each subject we computed the Pearson correlation between the upper triangular elements of the FC and SC matrixes. To assess whether such correlation varied between conditions we computed repeated Wilcoxon ranksum test across subjects and applied Bonferroni correction for multiple comparisons ($N^*(N-1)/2$ comparisons).

To investigate whether overall functional connectivity correlated with global structural connectivity across subjects we computed Pearson correlation between global values of FC and SC in each condition.

#### 6. Modelling whole-brain dynamics in patient groups and controls

For each condition we used the corresponding structural and functional connectivity data to build a generative whole-brain model capable of replicating the functional connectivity dynamics of the empirical conditions **(Fig. 1C).**

##### *The Hopf model*

The model consists of a set of differential equations with the following properties: (1) each equation describes the local dynamics of each node, (2) the equations are coupled via a structural connectivity matrix in such a way that each node's activity can influence and be influenced by other nodes' activity, (3) the equations contain a number of parameters that can be tuned for the collective behaviour of the model to mimic certain properties of the empirically collected data (for instance, the functional connectivity).

The connectivity between the different nodes is set to be equal to the empirical anatomical structural connectivity matrix $C = (C_{ij})$ derived from DSI (N= 115 areas), and the strength of these connections is scaled by the global coupling parameter G. This parameter is considered to reflect the effectiveness of conductivity of the structural connections. For each node i, the BOLD activity (corresponding to its local dynamics) was modelled using the normal form of a supercritical Hopf bifurcation with bifurcation parameter $a_i$. According to this model, a bifurcation occurs at $a_i = 0$ (edge of criticality), so that when $a_i$ assumes negative values (subcriticality) the activity of the node is described as noise, while when it assumes positive values (supercriticality), the behaviour becomes oscillatory with an intrinsic frequency determined by the parameter $f_i = \omega_i/2\pi$. In the current work, we set the frequency within the 0.04-0.07 Hz range, and we derived it from the empirical data, as given by the averaged peak frequency of the narrowband BOLD signals of each brain region.
The whole brain dynamics at node i can be described by this pair of coupled equations:

$$\frac{dx_i}{dt} = [a_i - x_i^2 - y_i^2]x_i - \omega_i y_i + \mathrm{G}\sum_{j=1}^{N} C_{ij}(x_j - x_i) + \beta\eta_i(t), \qquad [3]$$

$$\frac{dy_i}{dt} = [a_i - x_i^2 - y_i^2]y_i + \omega_i y_i + \mathrm{G}\sum_{j=1}^{N} C_{ij}(y_j - y_i) + \beta\eta_i(t). \qquad [4]$$

where the variable $x_i$ models the BOLD signal of area $i$, and $\eta_i(t)$ is additive Gaussian noise with standard deviation $\beta = 0.04$ [37]. The matrix $C_{ij}$ was normalised by a factor of 0.2, in order to prevent full synchronization of the model, in line with previous literature [69].

##### *Model fitting: estimation of local and global parameters in patient groups and controls*

An independent model was built separately for each condition using the corresponding group-averaged structural connectivity, i.e., the SC averaged across subjects within the group. Free parameters (the G coupling and the N bifurcation parameters $a_i$, one for each node) were allowed to vary in order to evaluate relevant properties of the data. We evaluated how the global properties of connectivity were homogeneously affected in each condition, i.e., all pairwise connections were scaled by the same parameter of global coupling G. Moreover, we evaluated how local dynamics were heterogeneously affected in the different conditions, i.e., different values of the bifurcation parameter $a_i$ were assigned to each node i. To find the optimal combination of the N+1 free parameters G and $a_i$ (for i=1, …, N), we used the following procedure.

We iteratively ran the model for coupling values G from 0 to 2.5 (in steps of 0.05). For each value of G, the bifurcation parameters were optimized based on the empirical power spectral density of the BOLD signals in each node. In particular, we iteratively ran the model, updating the set of bifurcation parameters $\{a_1, \ldots, a_N\}$ at each step as will be later described. For any value of $a$, simulated BOLD signals were filtered in the 0.04-0.25 Hz band and the power spectrum $PS_i(f)$ was calculated for each node (i). We then defined the proportion of power in the 0.04-0.07 Hz band as:

$$p_i = \frac{\int_{0.04}^{0.07} PS_i(f)df}{\int_{0.04}^{0.25} PS_i(f)df} \qquad [5]$$

All local bifurcation parameters were initially set to $a_i = 0$ and then updated until convergence by an iterative gradient descendent strategy, i.e.:

$$a_{i,new} = a_{i,old} \pm \zeta\left(p_i^{emp} - p_i^{sim}\right), \text{ for } i = 1, \ldots, N, \qquad [6]$$

with $\zeta = 0.1$. In the previous equation $p_i^{emp} - p_i^{sim}$ represents how much the simulated PSD in 0.04-0.07 Hz at node i deviates from the empirical one. When the $p_i^{emp} > p_i^{sim}$ , $a_i$ is increased to increase the oscillation to noise ratio at node i, whereas in the opposite case, $a_i$ is decreased to obtain smaller amplitude oscillations. Note that all $a_i$ values are updated in parallel in each optimization step.

As a result of the previous procedure, we obtained a set of optimized bifurcation parameters $\{a_1, \ldots, a_N\}$ for each value of G in the range from 0 to 2.5 (in steps of 0.05). To evaluate the goodness of fit of the model for each G value in this range we used two complementary objective metrics. The first metric was the functional connectivity (FC), which characterizes the spatial structure of temporal correlations between signals across the entire recording period. We quantified the FC fit as the Pearson correlation between the empirical and simulated FC matrices of each condition as a function of parameter G.

The second metric was the functional connectivity dynamics (FCD), which is based on the iFC(t) matrix defined above (see Eq. 1) and quantifies how much spatial patterns of phase coupling tend to repeat over time. For a single participant session, where T time points were collected, the corresponding FCD matrix is defined as a TxT symmetric matrix whose $(t_1, t_2)$ entry is defined by the cosine similarity between the upper triangular parts of the two matrices $iFC(t_1)$ and $iFC(t_2)$. At each value of G, we computed empirical and simulated FCD matrices, extracted their upper triangular elements and quantified the distance between the cumulative distribution functions of the two samples by computing the Kolmogorov-Smirnov (KS) statistics.

We ran 300 simulations for each value of G with their corresponding optimal set of bifurcation parameters and computed the mean FC fit and the mean KS distance across iterations. We considered the combination of G and $a$ parameter values where the mean KS distance was minimised as the optimal working point of the model. The optimal G and the corresponding optimal a (averaged across the 300 iterations) were selected for each condition independently.

#### 7. Additional statistical analyses

##### *Comparison of global coupling and local bifurcation parameters across conditions*

To disentangle global and local components of functional alterations in EP groups as compared to HC we separately compared the optimal G and the optimal $a$ parameter between conditions.

To evaluate the global change in functional connectivity we compared the optimal global coupling obtained in each EP group with that obtained in HC by applying a t-test test between optimal values of the parameter across repetitions. Results were corrected with Bonferroni analysis (N=3 comparisons) and a threshold was set at $\alpha$=0.05 to evaluate statistical significance.

To evaluate the change in local dynamics in each EP group as compared to HC we first computed the mean value across areas and then applied t-test between mean values of parameter $a$ across repetitions. Finally, we computed the difference between the mean and the median of the distribution of optimal $a$ to evaluate the influence of extreme values and compared it between conditions through unpaired t-test. Results were corrected with Bonferroni analysis and a threshold was set at $\alpha$ =0.05 to evaluate statistical significance.

##### *The role of functional and structural strength on each node's bifurcation parameter*

At this stage, we wanted to understand how the measured healthy structural and functional connectivity strengths in each node might relate to the corresponding optimal bifurcation parameters in the healthy model and to their alterations in the pathological condition. We made a general linear model of $a_i$ in HC ($a_i^{\mathrm{HC}}$, dependent variable) as a function of the functional and the structural node strengths, and their interactions ($FCS_i^{\mathrm{HC}}$, $SCS_i^{\mathrm{HC}}$, independent variables). Note that $FCS_i^{\mathrm{HC}}$ and $SCS_i^{\mathrm{HC}}$ are computed by averaging the functional and structural node strength, respectively, across subjects within the HC group. We compared the variance explained of this model with the variance explained of a model which included $FCS_i^{\mathrm{HC}}$ (or $SCS_i^{\mathrm{HC}}$, respectively) as the sole predictor. We then used the increase of variance explained as measure of the contribution of $SCS_i^{\mathrm{HC}}$ ($FCS_i^{\mathrm{HC}}$, respectively) in explaining the values of $a_i$.

We also investigated how the change in the optimal value of bifurcation parameters in each pathological condition as compared to HC ($\Delta a_i^{\mathrm{EP,HC}}$) relates with the corresponding values of $FCS_i^{\mathrm{HC}}$ *and* $SCS_i^{\mathrm{HC}}$. $\Delta a_i^{\mathrm{EP,HC}}$ was quantified as the difference between the absolute optimal value of $a$ in healthy controls minus the corresponding value in each subgroup of patients for that area. For completeness, the difference was computed in alternative ways as well, reported in supplementary materials (Supplementary Fig.3).

##### *Correlation between FC strength alterations and changes in the local bifurcation parameter*

To investigate whether alterations in the optimal value of the bifurcation parameter could underly changes in empirical functional connectivity in the pathological conditions, we explored the relation between these two variables. To this aim, we computed the Pearson correlation between the change in empirical functional connectivity in each group of patients ($\Delta FCS_i^{\mathrm{EP,HC}}$) and the corresponding change in the optimal value of bifurcation parameters ($\Delta a_i^{\mathrm{EP,HC}}$). $\Delta FCS_i^{\mathrm{EP,HC}}$ and $\Delta a_i^{\mathrm{EP,HC}}$ were quantified as described in the previous sections. A threshold was set at α=0.05 to evaluate statistical significance.

#### 8. The impact of local alterations on the network dynamics

As a following step, we decided to better investigate how local bifurcation parameters would affect global dynamical properties of the network. Based on equations 3 and 4 we hypothesised that negative values of the bifurcation parameter would have a damping effect on incoming oscillations, thereby decreasing the global synchronization and altering the spread of a signal across the network.

To test these hypotheses, we built a simplified network of coupled Hopf oscillators. Specifically, we built a network organised into two communities connected through a single node (hub). The strength of structural connection of each node was set to be equal to 0.12 toward the other nodes within the same community and equal to 0.01 toward all nodes within the other community. The strength of structural connection of the node that represented the hub of connectivity was set to be 0.2 with all nodes in the network. All parameters were selected to be compatible with measured ranges of empirical values. The global coupling was fixed to be equal to an intermediate value of G=0.3, previously found to be in the range of optimal values in real-case scenarios. The intrinsic frequency and the noise were fixed to be respectively equal to f=0.05 Hz and β=0.04 in all nodes, to be comparable with empirical data.

##### Spontaneous network dynamics and network response to external inputs

The bifurcation parameter was initially set to the point of criticality ($a = 0$) in all nodes. We then studied the network behaviour under two types of settings. On one hand, we changed the bifurcation parameter of the connectivity hub and/or of a weakly connected node towards increasingly negative values (subcriticality). In particular, the model was iteratively run for a range of $a$ going from $a = 0$ to $a = -3$ . In all cases, we studied the effect of this change on the spontaneous dynamics of the network, using the average FC as the main metric of interest. On the other hand, we also studied the effect of this change on the network response to external oscillatory inputs. In particular, we introduced an increasing number of synchronising stimuli applied to different nodes and measured the behaviour of the network under the perturbed dynamics.
In what follows, we describe the scenarios under which the model was tested (Fig. 4) to investigate different aspects of the impact of local bifurcation parameter on the network:

- Our first aim was to investigate the role of the local parameter $a$ in modulating the ability of a node to transmit incomings signals to the rest of the network and influencing its global synchronization. To this goal, for each value of $a$ (in the range from 0 to 3), we perturbed the hub at increasing intensity (Fig. 4A). More specifically we connected to the hub an increasing number (1 to 8) of external nodes oscillating synchronously at f = 0.05 Hz. We then analysed how it affected the synchronisation of the network by computing the global functional connectivity at each iteration.
- Then, we aimed at investigating how the local parameter $a$ influences the segregation of the network by affecting the spread of incoming synchronising stimuli between separated communities (Fig. 4B). To this aim, we perturbed a node in one of the two communities while changing the value of $a$ associated to the hub and measured the change in functional connectivity levels within the same and the other community.
- Finally, we investigated the influence on such network of a very disconnected node (Fig. 4C-E). We therefore included in the original network an additional node, whose strength of structural connectivity with the rest of the network was set to 0.01. We then investigated how changes in the local bifurcation parameter of the disconnected node and/or in the hub

influence the connectivity pattern of the network at rest, as well as its response to incoming stimuli.

### SUPPLEMENTARY RESULTS

#### 1. Additional analysis on group EP2

Stage2 patients were not included in the main analysis as they represent a more heterogeneous group of patients, including all subjects with full remission after the first episode that have not yet relapsed at the moment of the scan [ranging from 0.14 DOI (less than 2 months) to 4.61 DOI (4 years and almost 8 months)]. Therefore, both subjects fully recovered and subjects at a very early stage that may or may not relapse at any time after the scan are included in this group, complicating interpretation of the results. Nevertheless, for sake of completeness, we repeated the whole pipeline of analyses on stage 2 patients.

At the empirical level, no clear trend of alteration could be detected in EP2 patients in pairwise difference nor in strength **(Supplementary Fig. S6A, B)**. When mean strength difference was computed, EP2 patients showed a small reduction [-0.61 ± 0.02] (mean ± SE), $p<0.05$], but the effect size was much smaller (Cohen´s d=0.27) **(Supplementary Fig. S6C)**. Moreover, this difference was not replicated when computing functional connectivity as time averaged phase coherence **(Supplementary Fig. S2C)**.
No reduction was found in the optimal value of global coupling between HC and EP2 patients, where optimal resulted to be 1.32± 0.05 (mean ± SE across repetitions) (S**upplementary Fig. S6D**).
Regarding alterations in the local bifurcation parameters **(Supplementary Fig. S6E)**, EP2 patients did not exhibit the loss of heterogeneity and the shrink toward values around zero seen in EP3 patients. Instead, the alteration was in the opposite direction [-0.0042±3e-4] (grand mean ± SE across repetitions) **(Supplementary Fig. S6E)**. Similarly, in this subgroup neither the difference between the mean and the median across all ROIs nor the area below the curve were decreased compared to HC, but rather increased (**Supplementary Fig. S6F**).

### Supplementary Tables

**Supplementary Table 1: demographic data**

| | HC (n=128) | EPP (n=88) | stage II (n=37) | stage III (n=51) | stage IIIa (n=20) | stage IIIb (n=22) | stage IIIc (n=9) | EP3R (n=31) | EP3NR (n=20) |
|---|---|---|---|---|---|---|---|---|---|
| **Age** | 25,80 | 24,58 | 23,29 | 25,87 | 25,81 | 25,05 | 26,48 | 25,46 | 25,81 |
| **Handedness, %R** | 84,38% | 87,50% | 91,89% | 84,31% | 80,00% | 86,36% | 66,67% | 80,65% | 80,00% |
| **DOI (years)** | - | 1,91 | 0,81 | 3,02 | 2,24 | 1,99 | 4,91 | 2,83 | 2,24 |
| **CPZeq (mg/day)** | - | 349,51 | 336,98 | 362,03 | 372,96 | 351,05 | 390,24 | 362,43 | 372,96 |
| **PANSS positive** | - | 12,94 | 12,43 | 13,44 | 14,75 | 12,09 | 13,56 | 12,52 | 14,75 |
| **PANSS negative** | - | 16,00 | 15,34 | 16,66 | 18,05 | 14,68 | 17,56 | 15,52 | 18,05 |
| **PANSS general** | - | 32,83 | 32,97 | 32,68 | 36,75 | 31,09 | 30,11 | 30,81 | 36,75 |
| **PANSS total** | - | 61,76 | 60,74 | 62,78 | 69,55 | 57,86 | 61,22 | 58,84 | 69,55 |
| **GAF** | 83,84 | 56,79 | 57,71 | 55,88 | 50,45 | 58,73 | 57,44 | 58,35 | 50,45 |

**Supplementary Table 2: statistics on demographic data**

| | p-value HC/EPP | p-value II/III | p-value EP3R/EP3NR |
|---|---|---|---|
| **Age** | 0,13 | 0,03* | 0,69 |
| **Handedness, %R** | 0,35 | 0,04* | 0,72 |
| **Dur.Illness (years)** | - | <0,01* | 0,40 |
| **CPZeq (mg/day)** | - | 0,75 | 0,83 |
| **PANSS positive** | - | 0,34 | 0,08 |
| **PANSS negative** | - | 0,45 | 0,22 |
| **PANSS general** | - | 0,95 | 0,01* |
| **PANSS total** | - | 0,57 | 0,02* |
| **GAF** | <0,01* | 0,45 | 0,04* |

**Table3: Brain areas with extreme negative values in healthy controls**

| ***Optimal a* < -2MAD (-0.1548)** |
|---|
| R hipp presubic |
| R hipp ca4 |
| R hipp molecular layer |
| R vdc ventraldc |
| R hypothalamus |
| L entorhinal |
| L pallidum |
| L hipp presubic |
| L hipp ca4 |
| L hipp gcdg |
| L vdc ventraldc |
| L hypothalamus |
| brain-stem-midbrain |
| brain-stem-scp |

### Supplementary Figures

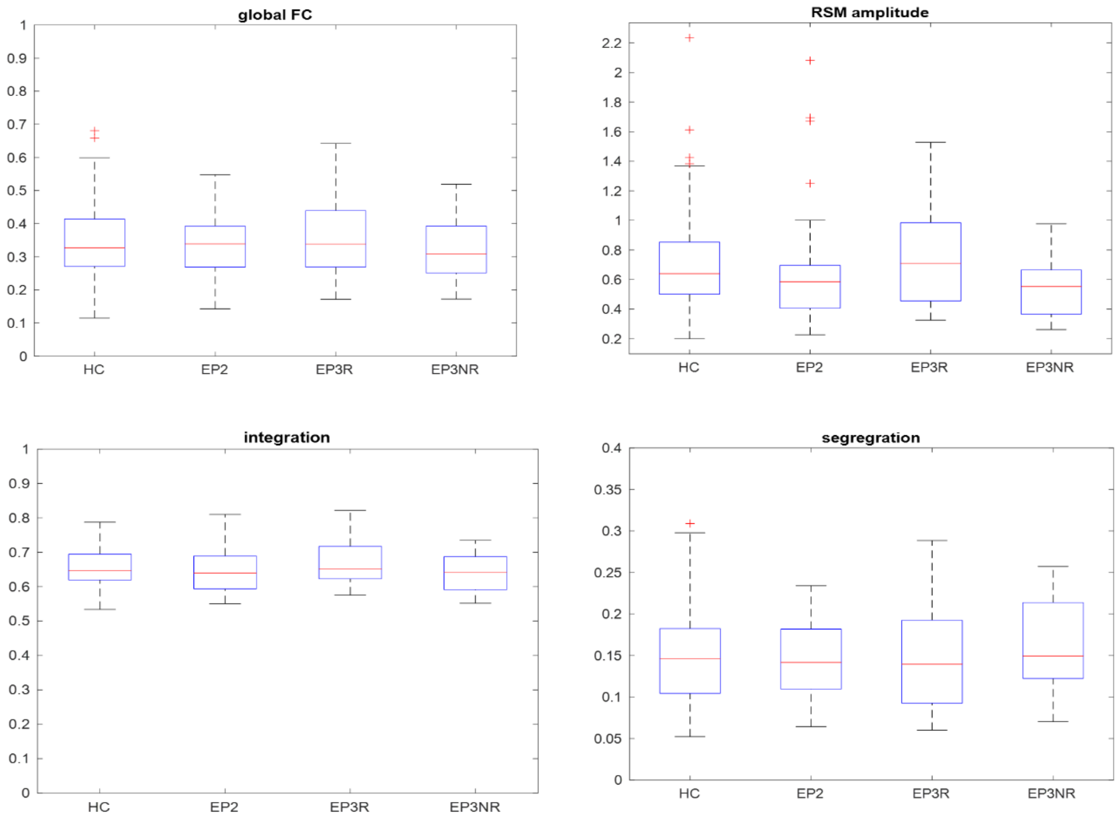

***Supplementary Fig.1: global e*mpirical FC measures.** *No significant difference between conditions was found when comparing the global functional connectivity (FC), the root-mean-square amplitude (RMS) amplitude, the integration and the segregation. Nevertheless, small but consistently opposite trends of alterations between the pathological groups as compared to healthy controls could be observed.*

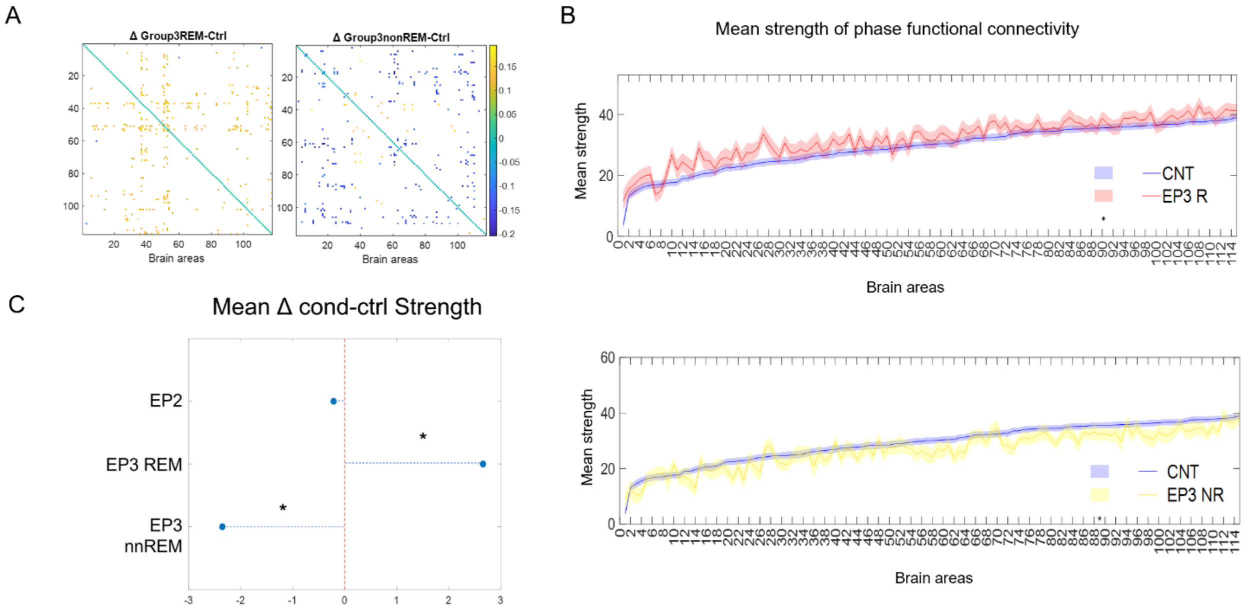

***Supplementary Fig.2: empirical phaseFC alterations.*** *To test the robustness of the results, we repeated all analysis computing FC with an alternative procedure to Pearson correlation. This procedure consists in measuring coupling between pair of nodes activity as the instantaneous phase difference between their signals at any time point (iFC) and only subsequently average it across time (phaseFC).* ***(A)*** *For each pair of brain areas difference in local functional connectivity in between the pathological conditions and the controls was computed. Only the areas with a significant difference (p<0.01) are showed, while all the other areas have been masked. No significance survived to multiple comparison analysis.* ***(B)*** *Group mean connectivity strength for each of the 115 regions, ordered by mean regional strength in healthy controls. Solid line and shaded area represent the median and the standard error across subjects. None of the comparisons survived multiple comparison analysis in any of the conditions. Both in (A) and (B) is possible to notice opposite trends of alteration in the two condition as compared to controls (increased FC in EP3R vs decreased FC in EPNR).* ***(C)*** *Mean strength difference across areas resulted significantly increased in EP3R in patients compared to healthy controls (p<0.001, Cohen effect=1.03), and significantly decreased in EP3NR patients (p <0.001, Cohen effect=0.75). No significant alteration was found in EP2 patients.*

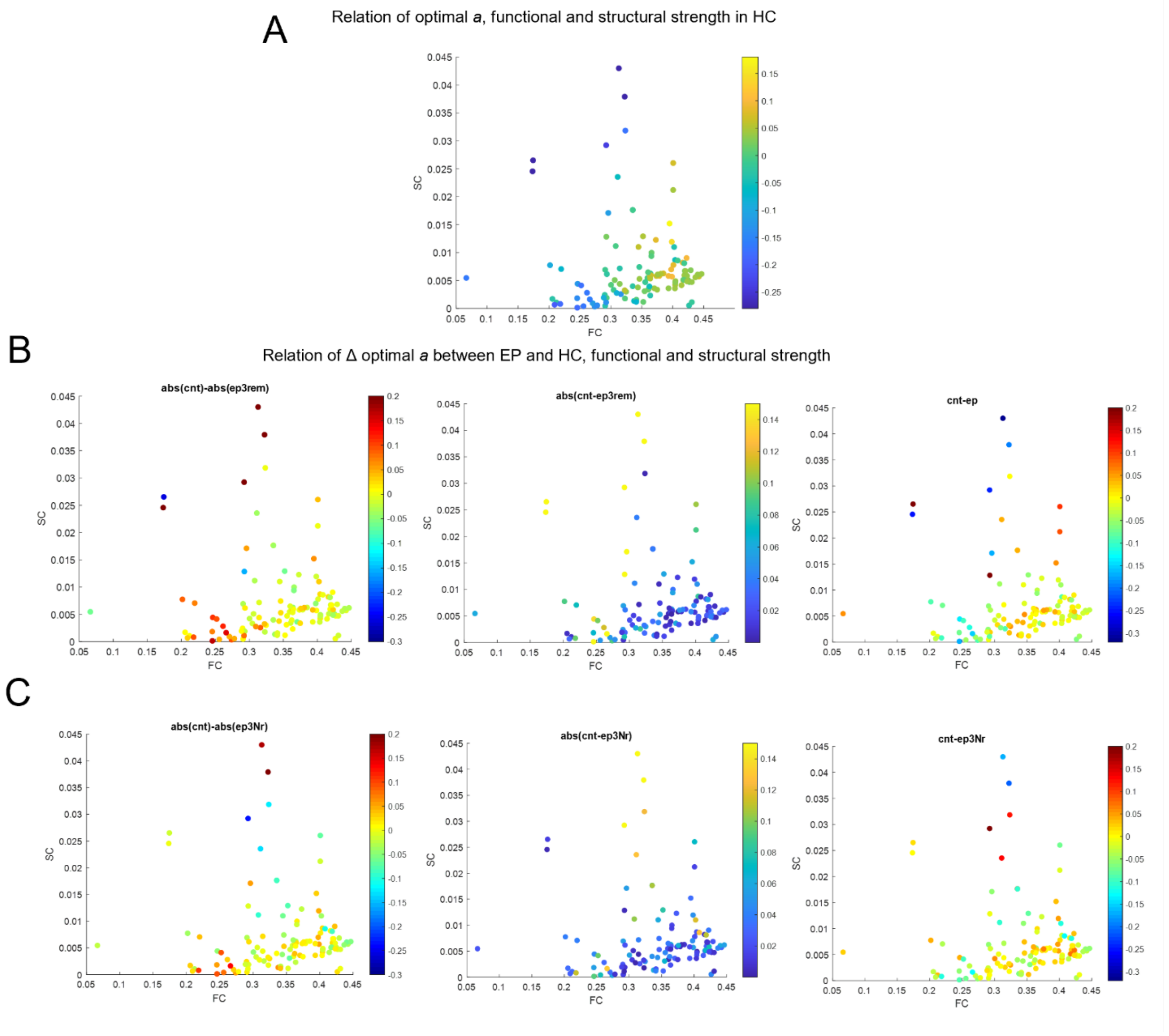

***Supplementary Fig.3: Relation of changes in optimal bifurcation parameters (Δa) with FC and SC strength.*** *We investigated how the change in the optimal value of bifurcation parameter in patients versus healthy controls (cnt) in each node ($\Delta a_i$) is related to its mean functional and structural strength ($FCS_i^{HC}, SCS_i^{HC}$).* ***(A)*** *Relation between mean optimal value of a in healthy controls (mean across repetitions), $FCS_i^{HC}$ and $SCS_i^{HC}$ (mean across subjects). The relation is illustrated in a 2D plot with SC and FC variables on the axes and values of optimal a represented as a gradient of colours (where blue indicates negative values and yellow positive values). Each dot represents one of the 115 areas.* ***(B)*** *Relation between mean $\Delta a_i$(mean across repetitions), $FCS_i^{HC}$ and $SCS_i^{HC}$ (mean across subjects) in remitting patients (EP3R). Since values of bifurcation parameter included both positive and negative value, and change could occur in both direction including change of sign, we computed Δa trough three different approaches to measure different dimension of the change. First, we computed the difference between absolute values between conditions (left panel), then we computed absolute value of the difference (central panel), and finally we computed the difference (right panel). The relation is illustrated in a 2D plot $FCS_i^{HC}$ and $SCS_i^{HC}$variables on the axes and values of optimal a represented as a gradient of colours (where blue indicates negative values, yellow indicate absence of change and red positive values). Each dot represents one of the 115 areas.* ***(C)*** *Relation between mean $\Delta a_i$ (across repetitions), $FCS_i^{HC}$ and $SCS_i^{HC}$ (mean across subjects) in remitting patients EP3NR).*

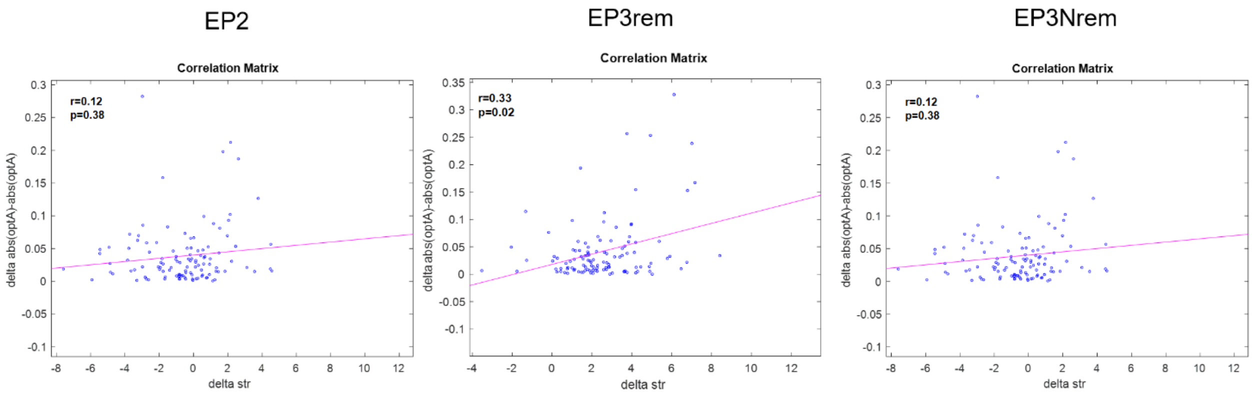

***Supplementary Fig.4: Correlation of changes in optimal bifurcation parameters (Δa) with changes in empirical FC strength ($\Delta FCS_i^{EP,HC}$).*** *We investigated whether the change in the optimal value of the bifurcation parameter ($\Delta a_i$) in each EP condition with respect to the HC correlated with the corresponding change in the empirical values of functional connectivity strength ($\Delta FCS_i^{EP,HC}$) illustrated in Fig.2. In EP3R (central panel), but not in the other two subgroups, we found a significant correlation between these two metrics, indicating that the alteration in the bifurcation parameter could be a relevant mechanism underlying the changes in connectivity.*

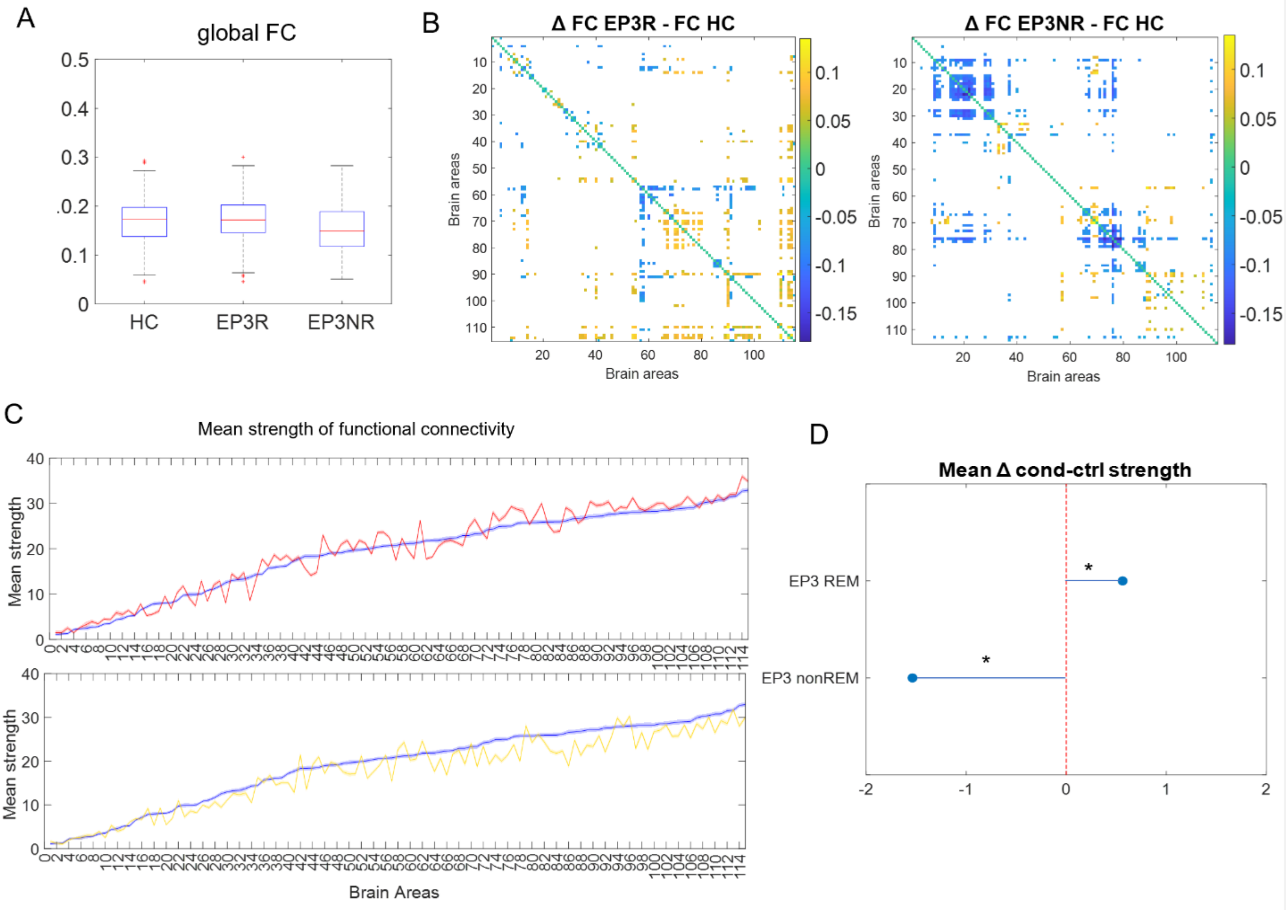

***Supplementary Fig.5: Comparisons in simulated FC measures between conditions:*** *empirical differences in FC measures between conditions were replicated in simulated data, further proving the goodness of the model in capturing relevant properties of the data. In particular in **(A)** As in empirical data, no significant difference between conditions was found when comparing the global functional connectivity, but we can notice that global FC tend to be lower in non-remittent patients than in remittent. **(B)** We can observe opposite directions of change in group mean pairwise functional connectivity between the simulated data of pathological conditions and of controls. Only the pair of brain areas with a significant difference with p<0.01 are shown, while all the other areas have been masked. **(C)** Group mean connectivity strength for each of the 115 regions, ordered by mean regional strength in simulated data of healthy controls. Solid lines and shaded areas represent the median and the standard error across repetitions(N=300), respectively. Different trends of alterations could be observed in the simulated data of two pathological groups as compared to healthy controls. **(D)** Mean strength difference across areas resulted significantly increased in simulated data of EP3R compared to simulated data of healthy controls, and significantly decreased in simulated data of EP3NR patients.*

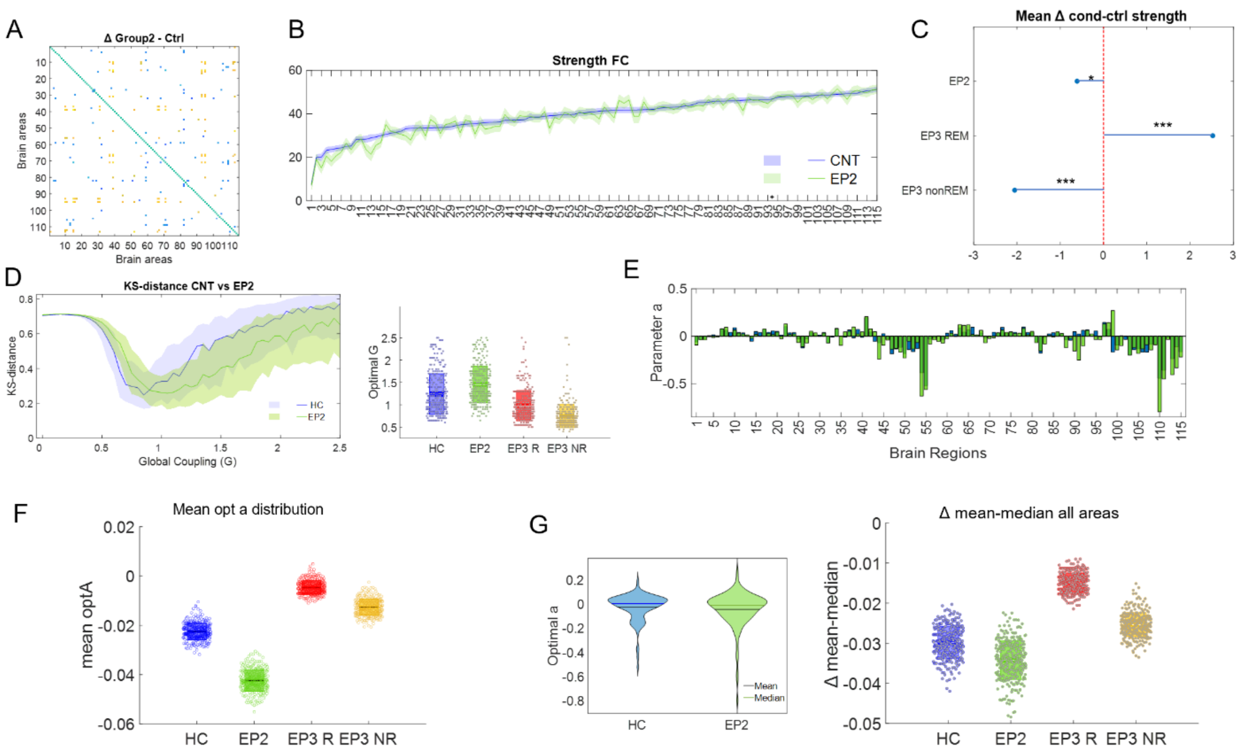

***Supplementary Fig.6: Analysis in EP2 patients**. In all panels, healthy controls are represented in blue, EP2 in green. **(A)** For each pair of brain areas difference in local functional connectivity in between the pathological conditions and the controls was computed. Only the areas with a significant difference (p<0.01) are showed, while all the other areas have been masked. **(B)** Group mean connectivity strength for each of the 115 regions, ordered by mean regional strength in healthy controls. Solid line and shaded area represent the median and the standard error across subjects. **(C)** Mean strength difference across areas resulted slightly decreased in EP2 patients as compared to controls. The difference is though not strongly relevant, as indicated by a small effect size (Cohen effect=0.27). **(D)** Fitting of free parameters in the model. The optimal combination of parameter was the one that minimised the Kolmogorov-Smirnov distance between the empirical and the model FCD distributions (left panel: solid line represents the median, shaded area represents the inter-quartile range across N=300 simulations).  Optimal global coupling G distribution (right panel). Each dot represents a simulation, and boxplots represent the mean of the measures' values with a 95% confidence interval (dark) and 1 SD (light). **(E)** Mean optimal value of a for each of the 115 nodes at the optimal value of G in healthy controls and patients. **(F)** Distribution of mean optimal bifurcations parameters across brain regions at optimal global coupling G. Each dot represents a simulation. Boxplots represent the mean of the measures' values with a 95% confidence interval (dark) and 1 SD (light). **(G)** Comparison of mean and median values of optimal bifurcation across brain regions between conditions. Violin plots on the left show optimal bifurcation parameters distribution across brain regions. Mean is plotted as a black line, while median is plotted as solid green line. Boxplots on the right represent the difference between mean and median in each condition. Each dot represents a simulation. Boxes represent the mean of the difference across simulations with a 95% confidence interval (dark) and 1 SD (light).*

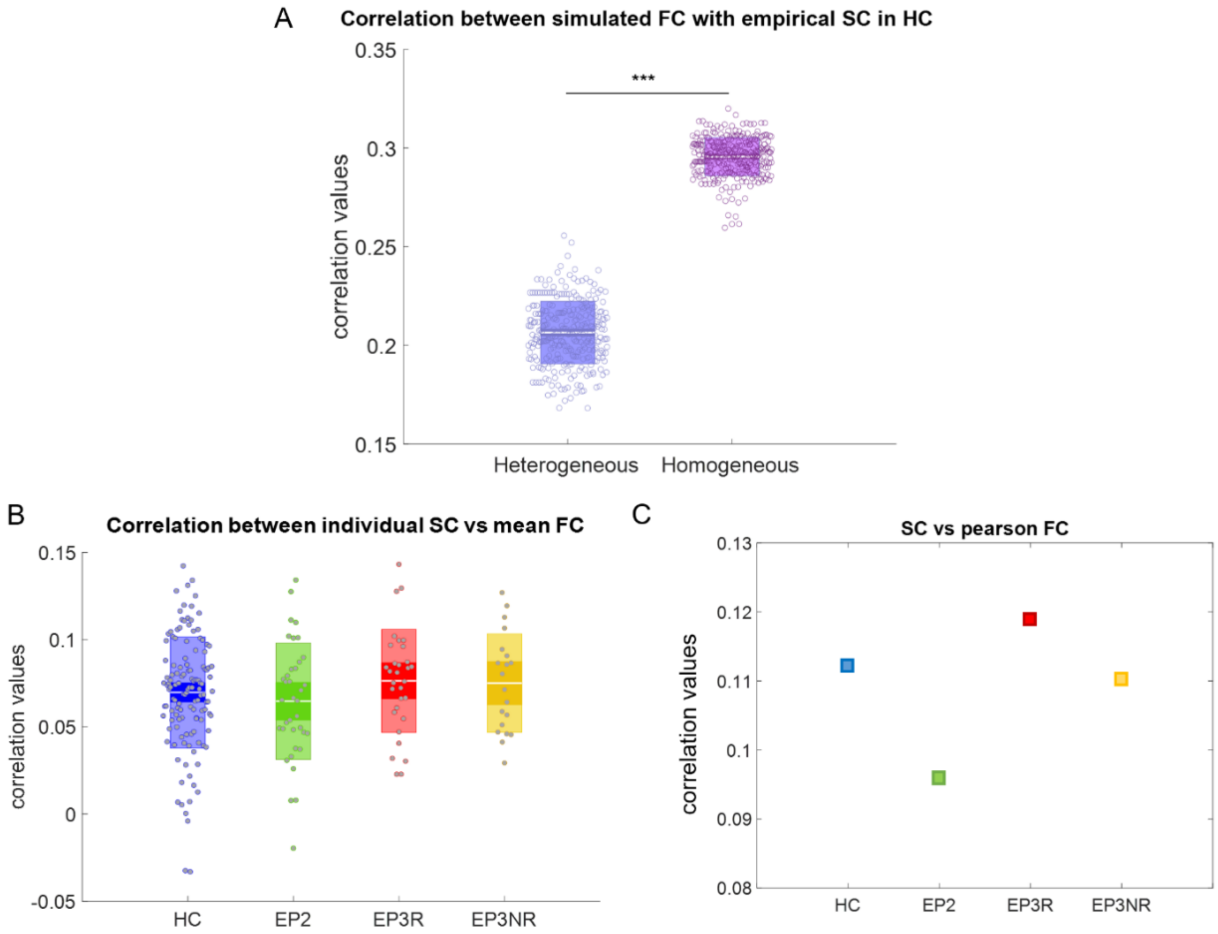

***Supplementary Fig.7: Structural-functional correlation. (A)*** *Correlation between simulated FC matrix and empirical SC of healthy controls in the homogeneous and heterogeneous model. We can see how the correlation is significantly decreased wen heterogeneity is included in the model, showing how it allows to escape structural constraints in the generation of functional pattern. (Reported from Fig. 3).* ***(B)*** *Correlation between individual empirical FC matrix and corresponding empirical SC of each subject in the different conditions.* ***(C)*** *Correlation between group average empirical FC matrix and empirical SC in each condition. In both* ***B*** *and* ***C*** *it is possible to notice ow the highest level of correlation correspond to the remitting patients, where the loss in heterogeneity of bifurcation parameters is biggest. At the same time, patients from group II, that exhibit increased heterogeneity in the model, are associated with a lower level of empirical functional-structural correlation.*
